# Immunogenicity and safety of a live-attenuated SARS-CoV-2 vaccine candidate based on multiple attenuation mechanisms

**DOI:** 10.1101/2024.03.03.582873

**Authors:** Mie Suzuki-Okutani, Shinya Okamura, Tanggis, Hitomi Sasaki, Suni Lee, Akiho Yoshida, Simon Goto, Mai Matsumoto, Mayuko Yamawaki, Toshiaki Miyazaki, Tatsuya Nakagawa, Masahito Ikawa, Wataru Kamitani, Shiro Takekawa, Koichi Yamanishi, Hirotaka Ebina

**Affiliations:** The Research Foundation for Microbial Diseases of Osaka University, Suita, Osaka, Japan; Virus Vaccine Group, BIKEN Innovative Vaccine Research Alliance Laboratories, Institute for Open and Transdisciplinary Research Initiatives, Osaka University, Suita, Osaka, Japan; Department of Experimental Genome Research, Research Institute for Microbial Diseases, Osaka University, Suita, Osaka, Japan; Center for Advanced Modalities and DDS (CAMaD), Osaka University, Suita, Osaka, Japan; Department of Infectious Diseases and Host Defense, Gunma University Graduate School of Medicine, Maebashi, Gunma, Japan; Virus Vaccine Group, BIKEN Innovative Vaccine Research Alliance Laboratories, Research institute for Microbial Diseases, Osaka University, Suita, Osaka, Japan; Center for Infectious Disease Education and Research (CiDER), Osaka University, Suita, Osaka, Japan

## Abstract

mRNA vaccines against SARS-CoV-2 were rapidly developed and effective during the pandemic. However, some limitations remain to be resolved, such as the short-lived induced immune response and certain adverse effects. Therefore, there is an urgent need to develop new vaccines that address these issues. While live-attenuated vaccines are a highly effective modality, they pose a risk of adverse effects, including virulence reversion. In the current study, we constructed a live-attenuated vaccine candidate, BK2102, combining naturally occurring virulence-attenuating mutations in the *NSP14*, *NSP1*, spike and *ORF7-8* coding regions. Intranasal inoculation with BK2102 induced humoral and cellular immune responses in Syrian hamsters without apparent tissue damage in the lungs, leading to protection against a SARS-CoV-2 D614G and an Omicron BA.5 strains. The neutralizing antibodies induced by BK2102 persisted for up to 364 days, which indicated that they confer long-term protection against infection. Furthermore, we confirmed the safety of BK2102 using transgenic (Tg) mice expressing human ACE2 (hACE2), that are highly susceptible to SARS-CoV-2. BK2102 did not kill the Tg mice, even when virus was administered at a dose of 10^6^ plaque-forming units (PFU), while 10^2^ PFU of the D614G strain or an attenuated strain lacking the furin cleavage site (FCS) of the spike was sufficient to kill mice. These results suggest that BK2102 is a promising live-vaccine candidate strain that confers long-term protection without significant virulence.

## Introduction

mRNA- and adenovirus vector-based vaccines have been successfully developed and used in clinical practice against SARS-CoV-2, the pathogen that caused the COVID-19 pandemic. These vaccines were highly effective, inducing robust humoral and cellular immunity (Baden et al., 2021; Polack et al., 2020). Nevertheless, certain drawbacks of SARS-CoV-2 vaccines remain to be addressed, including adverse effects, such as thrombosis, fever, and fatigue (Yasmin et al., 2023). Further, several boosters of the mRNA vaccines have been required to reactivate the immune response and increase efficacy against variants. New and improved vaccine modalities are therefore required for SARS-CoV-2 infection.

In general, live-attenuated vaccines are among the most effective vaccine modalities, as they induce humoral and cellular immunity, both systemically and locally (e.g., within mucosal surfaces), conferring protection against various infectious diseases (Hoft et al., 2017). For example, the live-attenuated poliovirus vaccine developed in 1962 was highly effective in reducing the spread of the disease (Sabin, 1985). Nevertheless, this modality has certain disadvantages, the most concerning being the risk of reversion to a virulent state as a result of mutations generated during viral replication *in vivo*. In fact, vaccine-derived paralytic polio was reported 38 years after the vaccine had been introduced, representing a threat to uninfected populations (Macklin et al., 2020). Advances in our basic knowledge of viruses have enabled the design of attenuated strains, such as that in the new type 2 oral polio vaccine (nOPV2), which is associated with a reduced risk of reversion due to genome modification (Yeh et al., 2020; Yeh et al., 2023). nOPV2 was recently approved by the World Health Organization, indicating that live-attenuated vaccines that overcome the risk of reversion are still in demand.

Various mechanisms leading to reduced pathogenicity of SARS-CoV-2 have been reported. A cold-adapted SARS-CoV-2 strain was isolated through passaging at low temperatures, eventually showing an attenuated phenotype (Seo and Jang, 2020). We also reported a temperature sensitive (TS) strain with low pathogenicity and sufficient immunogenicity *in vivo*, achieved through the introduction of multiple TS-related mutations (Yoshida et al., 2022). In addition to temperature sensitivity, several naturally occurring mutants exhibit attenuated phenotypes. Passaging SARS-CoV-2 in Vero cells facilitates isolation of strains with deletions in the furin cleavage site (FCS) (PRRAR) at the S1/S2 junction within the spike protein, resulting in low proliferation *in vivo* (Davidson et al., 2020; Peacock et al., 2021). Furthermore, a mutant that lacks four amino acids (PRRA) within the FCS did not induce weight loss in hamsters during challenge experiments (Hoffmann et al., 2020a; Hoffmann et al., 2020b; Johnson et al., 2021; Lau et al., 2020). Reportedly, S1/S2 cleavage is essential for TMPRSS2-mediated entry into host cells, and the deletion in the FCS is thought to reduce viral growth in lung cells, which express more TMPRSS2 than cells of the upper respiratory tract (Hoffmann *et al*., 2020a; Hoffmann *et al*., 2020b; Johnson *et al*., 2021; Lau *et al*., 2020). Furthermore, partial loss of *NSP1*, which correlates with a lower viral load and less severe symptoms of infection in SARS-CoV-2-infected patients, or of *ORF8*, which has been reported as associated with milder infection in humans, were also associated with attenuated phenotypes (Lin et al., 2021; Ueno et al., 2024; Young et al., 2020). Recovery of lost segments of viral genomes carrying deletions is more difficult than the reversion of amino acid substitutions (Bull, 2015). In this study, we designed and constructed live-attenuated vaccine candidates through a combination of substitutions and deletions involved in attenuation and reduced risk of virulent reversion. We then evaluated the candidate’s immunogenicity and safety in animal models, providing evidence that BK2102 is a promising vaccine candidate that confers prolonged protection.

## Results

### Construction of the live-attenuated vaccine candidate strains

Several genomic alterations are involved in the attenuation of SARS-CoV-2. In this study, we focused on deletions at three different sites within the viral genome: FCS within the spike protein, NSP1, and ORF7-8 (Supplementary Fig. 1A). While loss of the FCS inhibits virus-cell fusion mediated by TMPRSS2 in lung cells, partial deletion of NSP1 has been shown to impair viral proliferation *in vitro*, and the lack of ORF8 has been associated with milder symptoms and disease outcomes (Johnson *et al*., 2021; Lin *et al*., 2021; Ueno *et al*., 2024; Young *et al*., 2020; Zinzula, 2021). We previously obtained SARS-CoV-2 TS strains showing diverse attenuated phenotypes, and revealed that NSP3 L445F, NSP14 G248V, G416S, and A504V as well as NSP16 V67I were substitutions responsible for such phenotypes (Yoshida et al., 2022). Each substitution conferred some advantage for the development of an attenuated vaccine candidate with restricted proliferative capacity in deep regions of the body, such as lungs and brain. In addition, the presence of deletions is generally considered to confer a lower risk of reversion to a wild-type genotype when compared to amino acid substitutions. To this end, we constructed three candidates by combining several of the above-described mutations to design a safe live-attenuated vaccine strain (Supplementary Fig. 1A). The three candidates were inoculated locally into hamsters via the nasal route to mimic a natural infection. The immunogenicity of candidates 1 and 3 was much greater than that of candidate 2 (Supplementary Fig. 1B). Candidate 1, which has three deletions in the viral genome and contains three TS-responsible substitutions in NSP14, induced neutralizing antibodies when inoculated at a dose of 10^3^ PFU. Candidate 2, which has TS-related substitutions in both NSP3 and NSP14 and three deletions and, Candidate 3 which has only two deletions, were speculated to be excessively attenuated or to have a higher risk of virulent reversion. Taking these observations into consideration, we selected Candidate 1 for the vaccine, hereafter referred to as BK2102. The growth dynamics of BK2102 was evaluated at 32 °C and 37 °C (Fig. 1). It proliferated similarly to the wild-type B-1 strain at 32 °C (Fig. 1A), but the infectious virus titer was significantly lower compared to that of the wild type B-1 strain one day post-infection at 37 °C (Fig. 1B). Therefore, BK2102 showed a severe TS phenotype but could be amplified by incubating infected cells at 32 °C, which would also facilitate the manufacturing process.

**Fig. 1.**
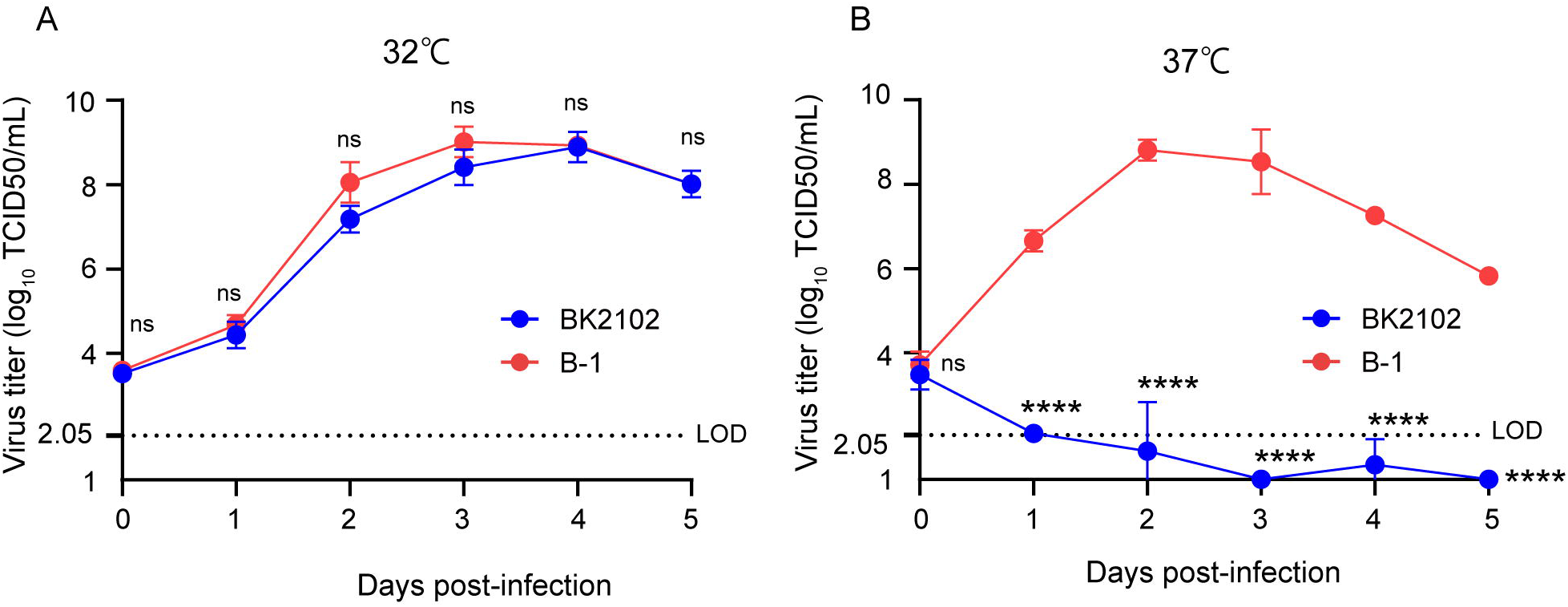
Growth dynamics of the vaccine candidate strain at different temperatures. Vero cells were infected with the wild-type parent B-1 (D614G) or the BK2102 vaccine candidate strains at a multiplicity of infection (MOI) = 0.01, and virus titers in the supernatants were determined for samples harvested every day, after incubating at 32 °C (A) or 37 °C (B). Infectious virus titers were determined using the TCID_50_ method. Symbols indicate the average of three independent experiments, and error bars represent the SD. The limit of detection (LOD) was 2.05 log10 TCID50/mL, and for samples below the LOD, the mean value was calculated as 1 log10 TCID50/mL. The dotted line represents the assay’s LOD. Days post-infection are indicated on the x-axis. For statistical analysis, two-way ANOVA with Sidak’s multiple comparison test was performed (ns, not significant; ****, *p* < 0.0001).

### BK2102 induced humoral and cellular immune responses

To evaluate immunogenicity, BK2102 was intranasally inoculated into Syrian hamsters (10^3^ and 10^4^ PFU/dose). Four weeks post-inoculation, spike-specific IgG was measured in the sera by ELISA (Fig. 2A), and the endpoint titers of the 10^3^ PFU- and the 10^4^ PFU-dose groups was 10^6.2^ and 10^6.1^, respectively. Neutralizing antibodies (Fig. 2B) against the D614G strain were detected in 9 of the 10 hamsters in each dose group, with titers ranging between 2^5^ and 2^9^ (Fig. 2B, left). Cross-reactivity of the neutralizing antibodies against the delta variant was also detected in 9 of 10 hamsters (titer range: 2^5^–2^8^) (Fig. 2B, middle) and 8 of 9 hamsters against gamma strain (Supplementary Fig. 5A), but that against the BA.5 variant was below the limit of detection in all hamsters (Fig. 2B, right). Furthermore, we performed BK2102 immunization of cynomolgus monkeys at a dose of 10^7^ PFU, and the serum neutralizing titer against the D614G strain was detected in two of the four monkeys (titer range: 2^4^–2^8^) (Supplementary Fig. 3A). Although a single dose did not raise neutralizing antibody titers in two of the four monkeys, three doses given at a two-week interval, induced neutralizing antibodies in all six monkeys (Supplementary Fig. 3B). The safety of BK2102 was also evaluated in these six monkeys, and no toxic effects were observed in any of the parameters assessed, including tissue damage, respiratory rate, functional observational battery (FOB), hematology, or fever (data not shown).

**Fig. 2.**
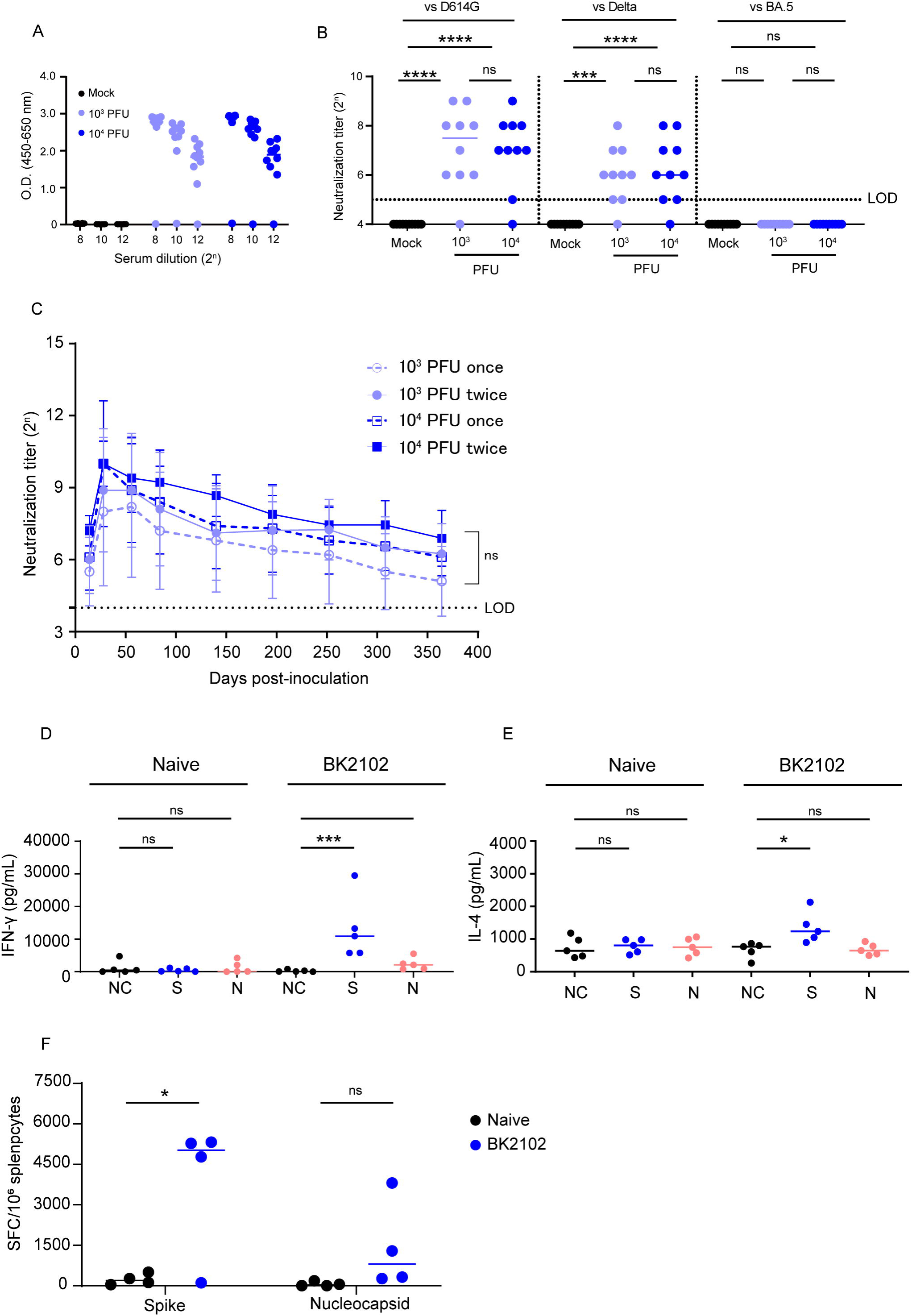
Immunogenicity of the vaccine candidate in hamsters. (A) Hamsters were inoculated with 1×10^3^ or 1×10^4^ PFU of BK2102 intranasally, and the serum was collected four weeks after inoculation. Spike-specific IgG in the sera of BK2102-inoculated hamsters and mock-treated hamsters was detected by ELISA. Symbols depict data of individual hamsters, and bars correspond to the median value. The limit of dilution is indicated in the x-axis. (B) Neutralizing antibodies in the sera were induced in BK2102-inoculated hamsters. Neutralizing antibodies in the sera were measured at day 28 post-inoculation using the following authentic SARS-CoV-2 strains: wild-type D614G (left), Delta (middle), and BA.5 (right). Symbols represent titers of individual animals, and the bars indicate the median. The LOD was 2^5^, and for samples below the LOD, the mean value was set to 2^4^. The dotted line represents the assay’s LOD. For statistical analysis, one-way ANOVA with Tukey’s multiple comparison test was performed (ns, not significant; ***, *p* < 0.001; ****, *p* < 0.0001). (C) Neutralizing antibodies persist in hamsters for at least 364 days. The neutralizing antibody titer against the authentic D614G wild-type strain was measured periodically in the sera of hamsters inoculated with BK2102 (once or twice at four-week intervals with 1×10^3^ or 1×10^4^ PFU) for about a year. Symbols represent the mean of 9 to 10 animals, and error bars represent the SD. The LOD was 2^4^, and for samples below the LOD, the mean value was set to 2^3^. The dotted line represents the assay’s LOD. For statistical analysis, two-way ANOVA with Tukey’s multiple comparison test was performed (ns, not significant). (D and E) Evaluation of the cellular immune response in BK2102-inoculated hamsters. Splenocytes were collected one week post-inoculation with 1×10^4^ PFU of BK2102 and were stimulated *in vitro* with spike or nucleocapsid peptide pools. IFN-γ (D) and IL-4 (E) in the supernatants were measured with commercially available ELISA kits (MABTECH AB and FineTest, respectively). Symbols depict data of individual hamsters, and bars indicate the median. For statistical analysis, one-way ANOVA with Tukey’s multiple comparison test was performed (ns, not significant; *, *p* < 0.05; ***, *p* < 0.001). (F) Evaluation of IFN-γ-secreting cells. Four hamsters were inoculated with 1×10^4^ PFU of BK2102 once and the splenocytes were collected a week later. Splenocytes were stimulated *in vitro* with spike or nucleocapsid peptide pools for 24 h. IFN-γ-secreting splenocytes were quantified by ELISPOT. Symbols depict data of individual hamsters, and bars indicate the median. For statistical analysis, two-way ANOVA with Sidak’s multiple comparison test was performed (ns, not significant; *, *p* < 0.05).

A short-lived immune response has been reported for current mRNA vaccines against SARS-CoV-2. For example, a reduction in neutralizing antibodies was observed in humans after six months (Zhang et al., 2022). In hamsters, these were undetectable after 250 days (Machado et al., 2023). Therefore, we evaluated the persistence of the immune response induced by BK2102 using a hamster model. We measured neutralizing antibody titers for up to 364 days after inoculation with BK2102. The titer peaks were observed 28 days after the first inoculation and slightly decreased, but were maintained until 364 days post-inoculation with a dose of 10^3^ or 10^4^ PFU. For example, the neutralizing antibody titer in the sera of hamsters inoculated with 10^3^ PFU was 2^8^ at 28 days and 2^5^ at 364 days post-inoculation. Two of the ten hamsters inoculated at a dose of 10^3^ PFU showed neutralizing antibody titers below the detection limit from the beginning, and those of another hamster in the same dose group began to decrease gradually from day 224 and fell below the detection limit on day 364 (Supplementary Fig. 4). However, the neutralizing antibody titers in hamsters inoculated with 10^4^ PFU did not exhibit such a decrease during the evaluation period. Remarkably, a single dose of BK2102 was sufficient to induce long-lasting immunity, and there was no need for booster immunization 28 days after the first inoculation (Fig. 2C).

Furthermore, we evaluated cellular immune responses following inoculation with BK2102. Antigen-specific IFN-γ and IL-4 production in spleen cells from inoculated hamsters was measured via ELISA after *in vitr*o re-stimulation with spike or nucleocapsid peptides. As shown in Figure 2D, spike peptide-specific IFN-γ production significantly increased in the splenocytes of BK2102-inoculated hamsters, as did the nucleocapsid peptide-specific IFN-γ production, although in this case it did not reach statistical significance. IL-4 production was significantly increased by spike-peptide stimulation (Fig. 2E). IFN-γ producing cells were also detected by enzyme-linked immunosorbent spot (ELISPOT) assays (Fig. 2F). In correlation with the ELISA results, significant induction of spike peptide-specific IFN-γ producing cells were detected in the splenocytes of BK2102-inoculated hamsters and the nucleocapsid peptide-specific IFN-γ-producing cells were also increased.

Moreover, a 10 μg-dose of a conventional mRNA vaccine prepared in-house, expressing spike protein of D614G strain of SARS-CoV-2, was intramuscularly injected into hamsters and compared to BK2102. Neutralizing antibody titers against the D614G strain showed no significant difference between the BK2102-inoculated group and the mRNA vaccine group (Supplementary Fig. 2A). Notably, under conditions that induced equal serum neutralizing antibody titers in hamsters, higher levels of spike-specific IgA in nasal wash samples were induced by BK2102 than by the conventional mRNA (Supplementary Fig. 2B). In addition, we qualitatively analyzed the spike-specific IgG subclasses (IgG1 and IgG2/3) in hamsters to evaluate the nature of the immune response induced by BK2102 (Supplementary Fig. 2C and D). When we inoculated BK2102 and our mRNA vaccine, total IgG antibodies were detected in both groups. IgG2/3 antibodies were detected in the sera of BK2102-inoculated hamsters, but we could not detect IgG1. On the other hand, mRNA-vaccinated hamsters showed both IgG subclasses (Supplementary Fig. 2C). As other studies have demonstrated that aluminum adjuvant preferentially induces a Th2 response (Marrack et al., 2009), we also administered recombinant spike protein with alum adjuvant as a control. The result was the same, since BK2102-inoculated hamsters showed only production of IgG2/3 (Supplementary Fig. 2D). IL-4 production and the presence of IgG1 reflects a Th2 response, while IFN-γ production and IgG2/3 are indicative of a Th1 response in hamsters (Kushawaha et al., 2011; Ploquin et al., 2013). Our results therefore suggest that BK2102 mainly induced a Th1 immune response in hamsters.

### BK2102 induced protective immunity against SARS-CoV-2 infection

Next, we performed challenge experiments with the SARS-CoV-2 D614G strain, BA.5 or gamma variants in order to evaluate whether the immune responses induced by BK2102 would protect against infection. All hamsters inoculated with BK2102 did not lose weight, whereas the naïve hamsters lost approximately 10% of their total body weight on day four or six post-challenge with the D614G or gamma strains (Fig. 3A and Supplementary Fig. 5B, respectively). When challenged with the BA.5 variant, all hamsters pre-inoculated with a dose of 10^4^ PFU of BK2102 and three of five hamsters pre-inoculated with a dose of 10^3^ PFU did not lose weight. However, the rest of the animals in the 10^3^ PFU dose group lost 5% of their total body weight, similarly to the naïve group (Fig. 3B).

**Fig. 3.**
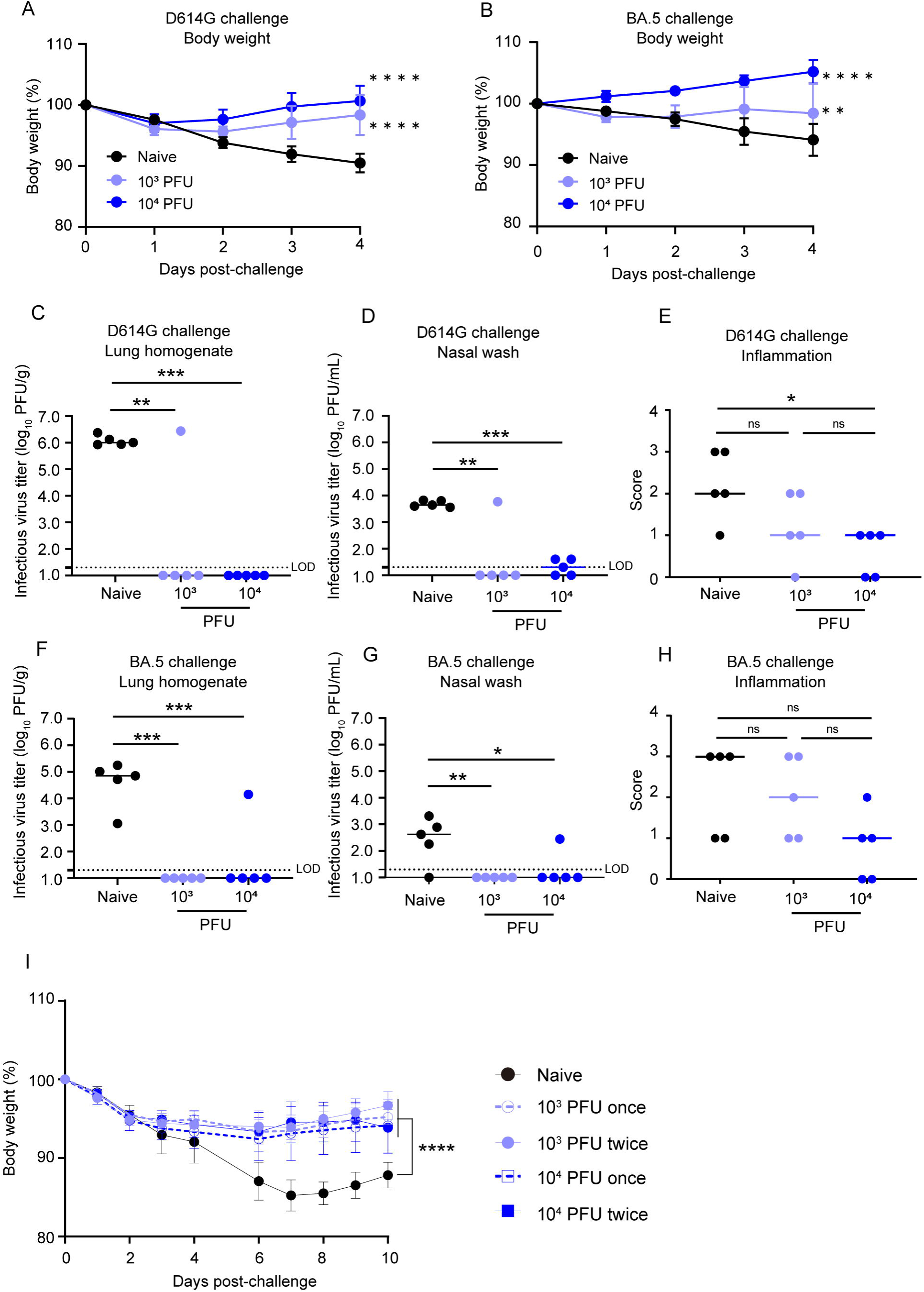
BK2102 induces protective immunity. (A and B) BK2102 protects hamsters against homologous and heterologous virus challenges. Hamsters that received a full vaccination protocol with the indicated doses of BK2102 were challenged with 3×10^5^ PFU of wild-type D614G (A) or BA.5 (B) strains, and their body weight was monitored for four days. Body weight is expressed as a percentage of the initial weight. Two-way ANOVA with Tukey’s multiple comparison test was performed for statistical analysis (**, *p* < 0.01; ****, *p* < 0.0001). (C, D, F and G) The infectious virus titer in the lungs and nasal wash specimens taken on day four post-challenge was measured via a plaque assay for the wild-type D614G strain (C and D) and for the BA.5 strain (F and G). The LOD was 1.3 log_10_ PFU/g or PFU/mL, and for samples below the LOD, the mean value was calculated as 1.0 log_10_ PFU/g or PFU/mL. The dotted line represents the assay’s LOD. One-way ANOVA with Dunnett’s multiple comparison test was performed for statistical analysis (ns, not significant; *, *p* < 0.05; **, *p* < 0.01; ***, *p* < 0.001). (E and H) Lung inflammation scores were determined via H&E staining of D614G-(E) and BA.5-challenged (H) hamsters. The percentage of the disrupted area in the entire visual field was classified as 0: not remarkable (< 10%); 1: minimal (10–50%); and 2: mild (50–70%). One-way ANOVA with Tukey’s multiple comparison test was performed for statistical analysis (ns, not significant; *, *p* < 0.05). (I) Weight changes after the challenge assay one year post-inoculation with BK2102. Hamsters inoculated with BK2102 were challenged with the wild-type D614G strain at 3×10^5^ PFU on 420 days. Nine-month-old elder hamsters were used as the naïve group. The symbols represent the average weight of the hamsters, and error bars indicate the mean SD. Two-way ANOVA with Tukey’s multiple comparison test was performed for statistical analysis (****, *p* < 0.0001).

In addition to body weight change, we determined infectious virus titers in lung homogenates and nasal wash specimens four days-post infection with the D614G strain or the BA.5 variant. The number of infectious viruses was significantly lower in hamsters inoculated with BK2102 than in naïve hamsters after challenge with the D614G strain (Fig. 3C and D). One hamster in the 10^3^ PFU dose group showed detectable levels of infectious virus following D614G challenge (Fig. 3C). This result was consistent with the undetectable levels of neutralizing antibodies in this animal (Fig. 2B). The virus titer in the lung and nasal wash of one animal in the 10^3^ PFU dose group was 6.4 log_10_ PFU/g and 3.8 log_10_ PFU/mL, respectively, and the mean virus titers in the naïve group was 6.1 log_10_ PFU/g and 3.7 log_10_ PFU/mL, respectively. Although the cross-reactivity of neutralizing antibodies against the BA.5 variant was below the limit of detection in all hamsters (Fig. 2B, right), no infectious virus was detected following challenge with the BA.5 variant in most of the vaccinated animals (Fig. 3F and G). The virus titer in the lung and nasal wash of one animal in 10^4^ PFU dose group was 4.2 log_10_ PFU/g and 2.5 log_10_ PFU/mL, respectively, and the mean virus titers in the naïve group which virus detected was 4.6 log_10_ PFU/g and 2.8 log_10_ PFU/mL, respectively. Lung tissue damage after the viral challenge was also evaluated in the hamsters. The inflammation score of hamsters inoculated with BK2102 was lower than that of naïve hamsters, regardless of the strain/variant used for the challenge (Fig. 3E and H, respectively). These results suggest that the immune response induced by BK2102 confers protection against infection that is not limited to the SARS-CoV-2 D614G strain, but also includes the BA.5 variant.

Furthermore, to evaluate whether the protection conferred after a full vaccination protocol would persist over time, hamsters with confirmed persistent immunity at day 364 post-inoculation (Fig. 2C) were challenged at day 420. In elderly naïve hamsters, a body weight loss of approximately 15% was observed seven days after infection with the D614G strain, whereas the BK2102-inoculated hamsters showed a lower weight decrease, with significant differences noted at this time point (Fig. 3I). Therefore, BK2102 induced a prolonged humoral immune response, which contributed to the protection against viral infection in hamsters.

We then evaluated whether BK2102 could inhibit onward transmission, as a previous report of a live-attenuated vaccine generated through codon-pair deoptimization (sCPD9) had suggested that an effective immune response within the nasal cavity would likely prevent it (Nouailles et al., 2023). The naïve group, the spike protein-inoculated (intra-muscularly) group, and the BK2102 intranasal inoculation groups were challenged with the SARS-CoV-2 D614G strain and co-housed with another group of naïve hamsters one day later (Supplementary Fig. 6A). Naïve animals co-housed with hamsters in the naïve or intramuscularly spike-alum vaccined groups showed slight weight loss. However, no weight loss was observed in hamsters co-housed with the hamsters that had been intranasally inoculated with BK2102 (Supplementary Fig. 6B). Therefore, intranasal inoculation of the BK2102 live-attenuated vaccine effectively prevented onward transmission, in line with a previous report (Nouailles *et al*., 2023).

### BK2102 caused localized tissue damage and conferred a low risk of transmission

Next, we evaluated the safety of BK2102 by assessing the tissue damage during acute infection. The lungs and whole heads of hamsters at day three post-infection were extracted and fixed with formalin (Fig. 4A). We evaluated inflammation and detected viral antigens in the lungs and multiple-depth sections of the nasal cavity (Fig. 4B, 4C and Supplementary Fig. 7). The D614G strain caused broad inflammation within the nasal cavity (from level 1 to 3) and lungs. Viral antigens were detected in the same areas, consistent with our previous report (Yoshida *et al*., 2022). However, in the BK2102-infected hamsters, viral antigens and weak-to-mild tissue damage were observed only in the anterior area of the nasal cavity, whereas no tissue damage or viral antigens were detected in the posterior area or lungs.

**Fig. 4.**
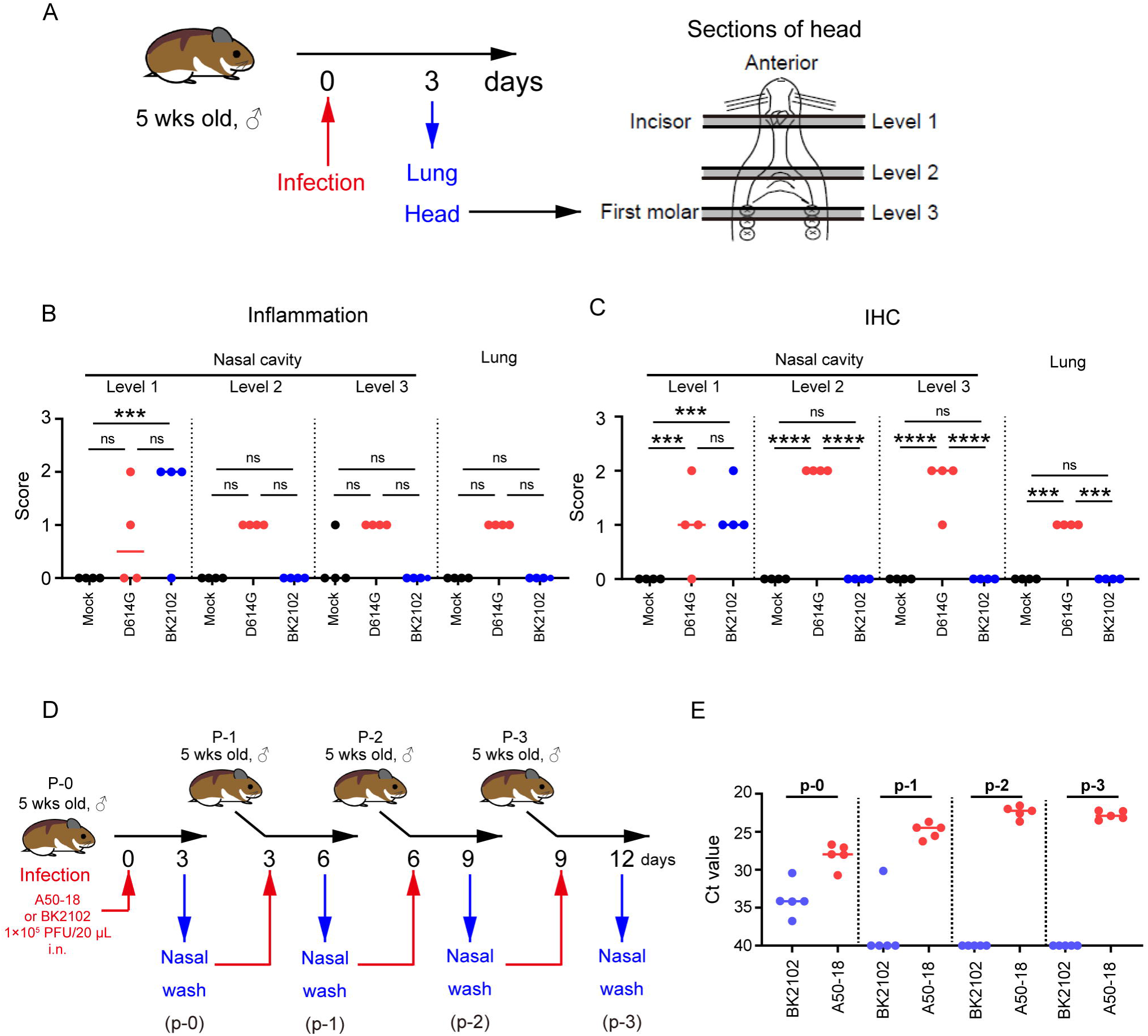
BK2102 caused localized tissue damage and posed as low risk of transmission. (A) Scheme for the evaluation of tissue damage in acute infection with BK2102 in a hamster model. The wild-type D614G strain was used as a positive control. (B) Inflammation score of nasal cavity sections and lungs determined via H&E. The percentage of the disrupted area in the entire visual field was classified as 0: not remarkable (< 10%); 1: minimal (10–50%); 2: mild (50–70%), respectively. One-way ANOVA with Tukey’s multiple comparison test was performed for statistical analysis (ns, not significant; ***, *p* < 0.001). (C) SARS-CoV-2 spike protein staining in the nasal cavity sections and lungs determined via immunohistochemistry using a SARS-CoV-2 spike RBD-specific antibody. The proportion of positive cells in the entire visual field was classified as 0: not remarkable (< 10%); 1: minimal (10–50%); and 2: mild (50–70%), respectively. One-way ANOVA with Tukey’s multiple comparison test was performed for statistical analysis (ns, not significant; ***, *p* < 0.001; ****, *p* < 0.0001). (D) Scheme for the evaluation of BK2102 transmission via *in vivo* passage in hamsters. The TS-strain A50-18 was used as a positive control. (E) Ct values obtained for the RT-PCR performed using RNA extracted from the nasal wash specimens.

The replication of BK2102 at the tip of the nasal cavity may facilitate transmission because infectious viruses are shed into the nasal fluid. In addition, virulent reversion may occur during replication *in vivo*. Therefore, we evaluated the risk of transmission and reversion to virulence by passaging *in vivo* using hamsters (Fig. 4D). SARS-CoV-2 A50-18 is a previously isolated TS strain, in which substitutions within the NSP14 protein alone account for the TS phenotype, without the need for deletions, such as those in NSP1, spike, or other accessory proteins (Yoshida et al., 2022). The viral genome was detected in all nasal wash specimens from A50-18 strain-infected hamsters during primary infection, and an increase in this amount was observed in subsequently passaged samples (Fig. 4E), which correlated with progressive weight loss (Supplementary Fig. 8). When we confirmed the sequence of the viruses detected in samples, we observed that the TS-responsible substitutions in NSP14 had reverted to the wild-type sequence (Table 3). The viral genome was detected in the nasal wash specimens from BK2102 infected hamsters during primary infection, but we could not detect it in the samples from subsequent hamsters, except for in one case in p-1 (Fig. 4E). In this individual animal, the viral genome was not detected in the samples from later passages. No weight loss was observed in any of the primary or subsequently infected hamsters, and we did not detect changes to the wild-type sequence (Table 3, Supplementary Fig. 8). Overall, our results suggest that BK2102 is a safe live-attenuated vaccine candidate with a low risk of virulent reversion.

**Table 1.** SARS-CoV-2 strains.

| **<u>REAGENT or RESOURCE</u>** | **<u>SOURCE</u>** | **<u>IDENTIFIER</u>** |
| --- | --- | --- |
| SARS-CoV-2: pre-alpha type, D614G, B-1 strain | Yoshida. et al.,2022 | NCBI: LC603286 |
| A50-18 strain | Yoshida. et al.,2022 | NCBI: LC603287 |
| L50-33 strain | Yoshida. et al.,2022 | NCBI: LC603289 |
| B-1 ΔFCS strain | Current study | N/A<sup>a</sup> |
| Candidate 1 (BK2102) | Current study | N/A<sup>a</sup> |
| Candidate 2 | Current study | N/A<sup>a</sup> |
| Candidate 3 | Current study | N/A<sup>a</sup> |
| SARS-CoV-2: delta variant, BK325 strain | Research Foundation for<br>Microbial Diseases of Osaka<br>University | N/A<sup>a</sup> |
| SARS-CoV-2: gamma variant, TY7-501 strain | National Institute of Infectious<br>Diseases | GISAID ID:<br>EPI_ISL_833366 |
| SARS-CoV-2: omicron variant, TY41-702 strain | National Institute of Infectious<br>Diseases | GISAID ID:<br>EPI_ISL_13241867 |

<sup>a</sup> Not applicable

**Table 2.**
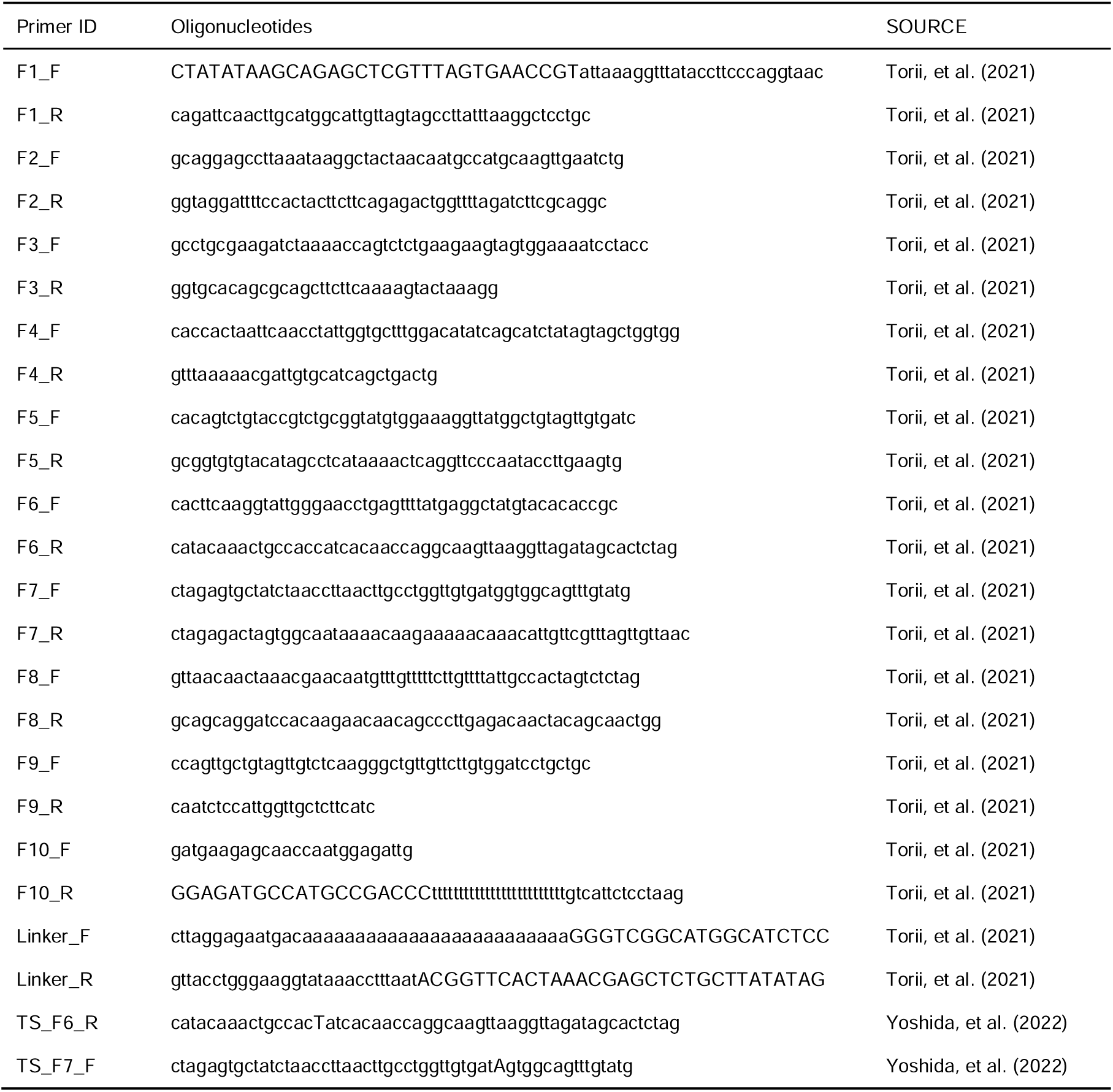
Primer list.

**Table 3.** Genetic variations of viruses passaged *in vivo*.

|  | Mutations |  |  |  |  |  |
| --- | --- | --- | --- | --- | --- | --- |
| | ΔG359 - A382 | G18782T | G19285A | C19550T | ΔA23598-G23624 | ΔC27549-T28251 |
|  | (NSP1) | (NSP14) | (NSP14) | (NSP14) | (Spike FCS) | (ORF7a-8) |
| A50-18 | N/A<sup>a</sup> | G | G/A | T | N/A<sup>a</sup> | N/A<sup>a</sup> |
|  | N/A<sup>a</sup> | G/T | G/A | T | N/A<sup>a</sup> | N/A<sup>a</sup> |
|  | N/A<sup>a</sup> | G/T | G/A | T | N/A<sup>a</sup> | N/A<sup>a</sup> |
|  | N/A<sup>a</sup> | G/T | G | T | N/A<sup>a</sup> | N/A<sup>a</sup> |
|  | N/A<sup>a</sup> | G/T | G/A | T | N/A<sup>a</sup> | N/A<sup>a</sup> |
| BK2102 | O<sup>b</sup> | T | A | T | O<sup>b</sup> | O<sup>b</sup> |
|  | O<sup>b</sup> | T | A | T | O<sup>b</sup> | O<sup>b</sup> |
|  | O<sup>b</sup> | T | A | T | O<sup>b</sup> | O<sup>b</sup> |
|  | O<sup>b</sup> | T | A | T | O<sup>b</sup> | O<sup>b</sup> |
|  | O<sup>b</sup> | T | A | T | O<sup>b</sup> | O<sup>b</sup> |

<sup>a</sup> Not applicable
<sup>b</sup> Same sequence as the inoculated virus

### BK2102 showed a favorable safety profile in Tg mice

hACE2 Tg mice are also used as animal models of SARS-CoV-2 infection (Asaka et al., 2021; Bao et al., 2020). We established a mouse line expressing hACE2 driven by the CAG promoter, and these mice were used to evaluate the safety of BK2102 live-attenuated vaccine candidate. hACE2 expression was detected not only in the respiratory tract, but also in various tissues such as the central nervous system, heart, skeletal muscle, digestive system (except the small intestine), spleen, and testis (Supplementary Fig. 9A). We evaluated the survival rates and body weight of Tg mice infected with various SARS-CoV-2 strains. All of the mice died after weight loss by infection with the D614G strain and even with the FCS deleted B-1 (B-1 ΔFCS) strain, previously established attenuated phenotype, within six days after receiving a dose of 10^2^ PFU (Fig. 5A and Supplementary Fig. 9B). Meanwhile, a higher survival rate was observed in mice infected with the L50-33 and A50-18 strains, which were previously isolated TS and live-attenuated strains (Yoshida *et al*., 2022), even at a dose of 10^5^ PFU. However, one mouse in each group infected with 10^4^ and 10^5^ PFU of the L50-33 strain died 10 days post-infection that is four days later than those in the D614G strain-infected group. This time lag before death was expected as the virus could have reverted during replication *in vivo*, being able to grow in deeper regions of the body. Thus, we evaluated the presence of infectious virus in the lungs and brains of mice that died following infection with the D614G, B-1 ΔFCS, and L50-33 strains. Infectious virus titers in the lungs were approximately 2.40 to 5.64 log_10_ PFU/g, while those in the brains were higher, at approximately 5.75 to 8.69 log_10_ PFU/g (Table 4). We noticed that mice exhibiting head nodding, intense running, jumping and repeated falling, died despite a generally mild inflammation in the lungs at necropsy (data not shown). These results suggest that Tg mice were killed due to replication of SARS-CoV-2 in the brain rather than in the lungs. Sanger sequencing analysis of viruses in the lungs and brains of mice that died following infection with the L50-33 strain revealed that the 445F substitution in NSP3, responsible for the TS phenotype of this strain, had reverted to the wild-type amino acid leucine (TTT → TTG), which is the same in the D614G and B-1 ΔFCS strains (CTT, Table 4). These results suggest that the mice died due to viral proliferation in the brain, where a small virus population lost its temperature sensitivity, becoming virulent. This mouse is a highly susceptible model of SARS-CoV-2 virus infection able to detect a few temperature-sensitive revertant viruses. In contrast to L50-33 with the NSP3-based TS phenotype, A50-18, harboring three TS-responsible substitutions in NSP14, did not kill any mice. Moreover, in the case of BK2102, no mice died or lost weight following infection, even at a dose of 10^6^ PFU (Fig. 5B and Supplementary Fig. 9C, respectively). Therefore, BK2102 is considered to have a low risk of virulent reversion, thus representing a suitable candidate for a safe live-attenuated vaccine.

**Fig. 5.**
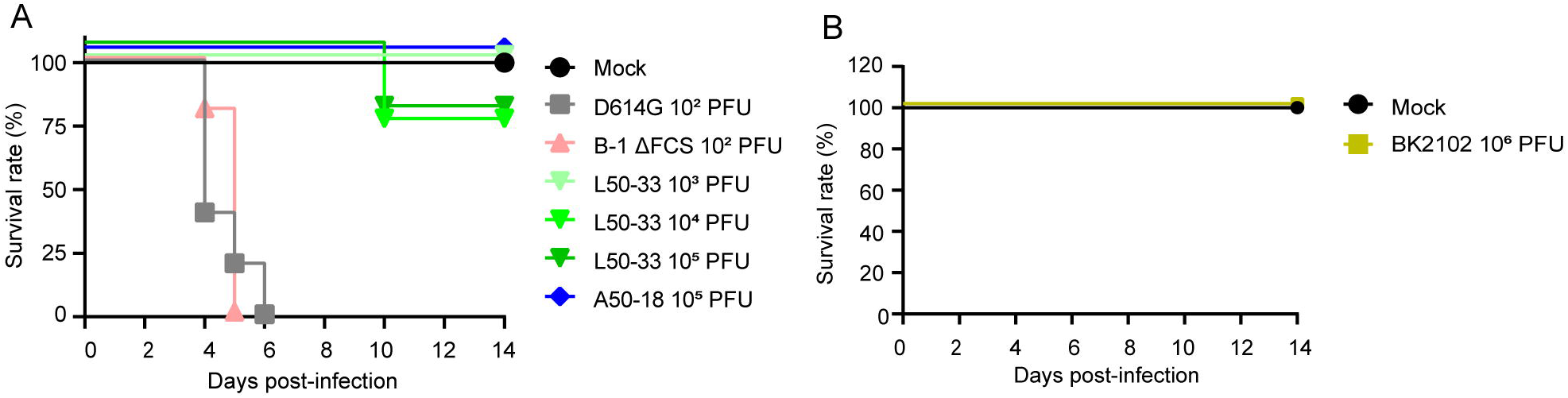
BK2102 showed a highly safe phenotype in Tg mice. (A and B) Survival rate of Tg mice infected with the wild-type D614G, B-1 ΔFCS, L50-33, and A50-18 TS strains (A) and BK2102 (B).

**Table 4.** NSP3 genetic variations in viruses recovered from infected Tg mice.

|  | n | Virus titer<br>(Log10 PFU/g) |  | TS substitution |
| --- | --- | --- | --- | --- |
|  |  | Brain | Lung | NSP3 |
| **D614G**<br>**(10<sup>2</sup> PFU infected)** | 5 | 6.94 ± 0.44<sup>a</sup> | 3.67 ± 0.26<sup>a</sup> | N/A<sup>b</sup> |
| **B-1 Δ FCS**<br>**(10<sup>2</sup> PFU infected)** | 5 | 8.69 ± 0.33<sup>a</sup> | 5.64 ± 0.62<sup>a</sup> | N/A<sup>b</sup> |
| **L50-33**<br>**(10<sup>4</sup> PFU infected)** | 1 | 5.75 | < 2.40 | F445L<br>(TTT → TTG) |
| **L50-33**<br>**(10<sup>5</sup> PFU infected)** | 1 | 7.24 | 2.40 | F445L<br>(TTT → TTG) |

<sup>a</sup> Average ± SD
<sup>b</sup> Not applicable

## Discussion

In this study, we evaluated candidates for a new live-attenuated SARS-CoV-2 vaccine. The candidate harboring more mutations exhibited lower immunogenicity than those with fewer mutations (Supplementary Fig. 1A), presumably, because excessive mutations hinder viral replication in the body, resulting in weaker immune responses. Deletions within viral genes are less prone to reversion to the wild-type genotype, and TS-associated substitutions restrict viral dissemination to deep or warmer regions of the body, including the brain, which makes them an attractive backbone for developing safe live-attenuated vaccines. Our BK2102 vaccine candidate was designed with three deletions in addition to TS-associated substitutions.

We previously reported that TS strains replicate only in the anterior regions of the nasal cavity and do not proliferate in the posterior areas or lungs of hamsters. However, these isolated TS strains posed a risk of reversion to virulence (Yoshida et al., 2022). K18-hACE2 Tg mice have been shown to die after infection with several SARS-CoV-2 strains due to viral proliferation in the brain (Natekar et al., 2022). To better assess the safety of BK2102, we generated and used CAG-hACE2 Tg mice in addition to hamsters. The TS strains (L50-33 and A50-18) showed an attenuated phenotype in this mouse model, emphasizing the ability of TS to limit viral replication and restrict replication-permissive regions by temperature. The TS-responsible substitutions in NSP14 were more stable than those in NSP3, as L50-33 caused the death of one mouse in each of the dose groups tested, whereas A50-18 did not. Additionally, infectious viruses lacking TS mutations were detected in the central nervous system and respiratory tracts of mice that died following L50-33 infection (Table 4). These findings suggest that our mouse model is particularly suitable for assessing safety and evaluating TS strains’ entry into the nervous system, as it enables the detection of even a small population of revertant viruses. This animal model also addresses regarding SARS-CoV-2 infection via the intranasal route causing central nervous system damage (Jha et al., 2021; Kumar et al., 2020).

BK2102, did not proliferate in the brains of Tg mice even at a dose of 10^6^ PFU, which was 10,000 times higher than the dose that killed mice infected with the D614G or B-1 ΔFCS strains (Fig. 5A and 5B). Importantly, BK2102 was not detected during passaging *in vivo* using naïve hamsters, probably due to the multiple defective mutations that controlled replication in the host animal, preventing sufficient proliferation for transmission. In contrast, the A50-18 strain, with a genetic background similar to BK2102 but lacking the three deletion mutations, showed an attenuated phenotype during primary infection, but reverted to virulence during replication in hamsters, leading to transmission of the virulent strain to naive animals (Fig. 4E). We hypothesized that even if TS-related substitutions are lost during primary infection, the remaining deletion mutations in BK2102 ensure low replication efficiency, preventing transmission. Indeed, mutations in the FCS coding region has been reported to reduce proliferation (Lau *et al*., 2020; Sasaki et al., 2021; Wang et al., 2021a) and to prevent transmission *in vivo* (Peacock *et al*., 2021).

Our vaccine candidate BK2102 induced humoral and cellular immune responses in hamsters, and animals were protected against challenge with the heterologous BA.5 variant, even though the neutralizing titer in serum was below the limit of detection (Fig. 2B, right). The nucleocapsid proteins of many coronaviruses are highly immunogenic and are abundantly expressed in infected cells, making them effective targets for antigen-specific T cells (Cong et al., 2020; Dutta et al., 2020; Hasanpourghadi et al., 2023). Studies have also reported the benefit of vaccination with the nucleocapsid protein of SARS-CoV-2, showing protective immunity in animal models vaccinated with the nucleocapsid protein alone (Primard et al., 2023) or in combination with the spike protein (Chiuppesi et al., 2022; Hasanpourghadi et al., 2023). In our hamster study, we observed cellular immune responses against both the nucleocapsid and spike protein. BK2102 may induce a cellular immune response against various structural proteins of SARS-CoV-2, providing protection against multiple variants. We also considered the potential induction of mucosal immunity. In this study, BK2102 induced spike-specific IgA in nasal wash (Supplementary Fig. 2B). Live-attenuated influenza vaccines have been reported to be effective against a broad range of variants due to the robust humoral and cellular immune responses they elicit, even in mucosal tissues (Thwaites et al., 2023). BK2102 was administered intranasally, and inhibition of onward transmission was observed (Supplementary Fig. 6), similar to what was reported for the SARS-CoV-2 live-attenuated vaccine candidate sCPD9 (Nouailles et al., 2023). Thus, we assumed that BK2102 also induced a mucosal immune response in addition to systemic humoral and cellular immunity, contributing to protection against mutant strains and prevention of onward transmission to other animals.

The findings of this study should be interpreted in light of certain limitations. First, the limited availability of analytical reagents for hamster models restricted the detailed immunological characterization of the response. Additionally, it took time to gather preclinical data due to the space-related restrictions of BSL3 facilities, which delayed the clinical trials for BK2102 until many individuals had already acquired immunity against SARS-CoV-2. It remains to be seen whether our candidate will be optimal for human use, as the immunogenicity of live-attenuated vaccines is generally influenced by pre-existing immunity. Finally, species-related differences in susceptibility must also be considered. The minimum infectious titer of SARS-CoV-2 has been reported as 10 TCID50 in hamsters and humans (Lindeboom et al., 2024; Rosenke et al., 2020), but 3.84×10^4^ PFU in monkeys(Johnston et al., 2021). When we inoculated this candidate into monkeys, they were less susceptible than hamsters, requiring a very high titer and multiple doses to induce immunity. Therefore, the dosage for first-in-human trials should be carefully optimized. It is also important to note that live-attenuated vaccines are contraindicated in immunosuppressed individuals or those with chronic diseases, and BK2102 is not intended for these populations.

The key to developing live-attenuated vaccines lies in balancing immunogenicity and safety. Most live-attenuated vaccine candidates, such as sCPD9 and CoviLiv^TM^ (which is in phase III clinical trial), have been evaluated with a primary focus on immunogenicity in animal models (Wang et al., 2021b). In this study, we rigorously assessed not only the immunogenicity, but also the safety of BK2102, demonstrating its superior safety profile. For example, sCPD9 and CoviLiv^TM^ are attenuated by codon deoptimization or a combination of codon deoptimization and FCS deletion (Trimpert et al., 2021; Wang *et al*., 2021b). These strategies affect viral proliferation but not necessarily virulence. The TS-responsible substitutions in NSP14 included in BK2102 selectively restrict the infection site, reducing the likelihood of lung and brain infection and enhancing safety.

Our findings also indicated that the TS substitutions in BK2102 made it difficult to passage the virus *in vivo*, limiting its spread to the central nervous system. Although attenuated strains with amino acid substitutions pose a risk of reversion to virulence, combining multiple modifications related to diverse attenuated phenotypes may allow for the construction of safer live-attenuated vaccine candidates. Among these modifications, deletions are particularly useful in reducing the risk of reversion to virulence, and TS-responsible substitutions significantly limit viral replication in deep regions of the body. This strategy of combining multiple modifications could be effective for the development of live-attenuated vaccines against other viruses, such as the Japanese encephalitis and influenza viruses, which have been reported to replicate in the brain and lung (Desai et al., 1995; Weinheimer et al., 2012).

## Material and methods

### Cells and viruses

Vero cells were purchased from ATCC and maintained in D-MEM supplemented with 10% FBS, penicillin (100 U/mL), and streptomycin (0.1 mg/mL). VeroE6/TMPRSS2 cells were obtained from the Japanese Collection of Research Bioresources (JCRB) cell bank and cultured in D-MEM supplemented with 10% FBS, penicillin (100 U/mL), streptomycin (0.1 mg/mL), and G-418 (1 mg/mL). We previously constructed baby hamster kidney (BHK) cells constitutively expressing the human angiotensin-converting enzyme 2 (hACE2) (BHK/hACE2 cells) (Okamura et al., 2023), and these cells were maintained in MEM, supplemented with 10% FBS, penicillin-streptomycin, and puromycin (3 μg/mL). The SARS-CoV-2 strains used in this study are listed in Table 1. The SARS-CoV-2 B-1 (D614G) strain was isolated from a clinical specimen, and TS derivative strains were obtained through random mutagenesis of this clinical isolate, as previously reported (Yoshida et al., 2022). SARS-CoV-2 delta and omicron variant were obtained from the Research Foundation for Microbial Diseases of Osaka University and the National Institute of Infectious Disease of Japan, respectively.

### Construction of viruses through circular polymerase extension reaction (CPER)

SARS-CoV-2 live-attenuated vaccine candidate strains and B-1 ΔFCS strain has genetic backgrounds similar to that of B-1 strain, in combination with following naturally occurring virulence-attenuating mutations. The mutations in the ORF7a-8, *NSP3*, *NSP14*, and *NSP16* coding regions are the same as those in the genomes of temperature-sensitive virus strains (L50-33, A50-18 and H50-11), described in Yoshida et al., 2022. The mutations in the spike FCS (_679_NSPRRARSV_687_ → I) and the NSP1 (_32_GDSVEEVL_39_) are the same as those in the genomes of a laboratory strain described in Davidson *et al.,*2020 and of a clinical isolate (accession: LC521925), respectively.

For construction of the strains we used the CPER method, in which 11 PCR-generated cDNA fragments covering the viral full genome plus another DNA fragment with controlling sequences are stitched into a circular DNA that can produce viral genomic RNA upon introduction to cells (Okamura et al., 2023; Torii et al., 2021). Primers used to prepare PCR fragments are listed in Table 2. The PCR and CPER reaction were performed using PrimeSTAR GXL DNA polymerase (Takara Bio, Cat# R050A). The CPER product was transfected into BHK/hACE2 cells using Lipofectamine LTX (Thermo Fisher Scientific). The cells were incubated at 32 °C in a CO_2_ incubator for one week. At this point, the supernatants were collected and transferred to 6-well plates which had been pre-seeded with VeroE6/TMPRSS2 cells. The viruses contained in the supernatants of these cells were collected when cytopathic effects (CPEs) were clearly observed.

### Virus titration

The infectious titer of SARS-CoV-2 was determined based on the median tissue culture infectious dose (TCID_50_) or plaque formation assay (PFA). In order to obtain the TCID_50_, virus-containing samples were serially diluted with D-MEM supplemented with 2% FBS. Fifty μL of each diluted sample were used to infect Vero cells in 96-well plates. The cells were fixed with 10% formalin after incubating at 32 °C for four days and stained with crystal violet solution. The TCID_50_ was calculated using the Behrens-Karber method. For the calculation of PFU, 500 μL of the diluted samples were added to confluent Vero cells in 6-well plates and incubated at 32 °C for 3 h to allow virus adsorption. The supernatant was removed, and the cells were washed with D-PBS. Subsequently, 2 mL of 1% SeaPlaque agarose dissolved in D-MEM and supplemented with 2% FBS were layered on the cells. The plates were incubated at 32 °C in a CO_2_ incubator for three days, fixed with 10% formalin, and stained with crystal violet solution. Visible plaques were counted to calculate PFU.

### Viral proliferation assay at various temperatures

The SARS-CoV-2 B-1 (D614G) strain or vaccine candidate strain was used to infect Vero cells at a multiplicity of infection (MOI) of 0.01. Infected cells were cultured at 32 °C or 37 °C, and a part of the supernatant was collected daily and stored at -80 °C. The viral titers of these samples were determined using the TCID_50_ assay described above. Each experiment was performed in triplicates.

### Quantitative RT-PCR

Viral RNA was extracted from various samples using the IndiSpin Pathogen Kit (INDICAL BIOSCIENCE) following the manufacturer’s instructions. RNA was quantified using a DetectAmp SARS-CoV-2 RT-PCR kit (Sysmex). Five µL of a 100-fold dilution of the extracted RNA were used to perform the reverse transcription and subsequent qPCR reactions in a 7500 Fast Real-time PCR System (Applied Biosystems). The reactions were performed in triplicate.

### Neutralization assay

The neutralizing activity of sera was evaluated using authentic SARS-CoV-2. One hundred PFU of authentic virus were mixed with serially diluted serum samples and incubated at 37 °C for 1 h. The mixtures were then transferred to confluent Vero cells in 96-well plates. The infected cells were fixed with 10% formalin and stained with crystal violet solution after an incubation period of three days at 32 °C. Neutralization titers were calculated as the inverse of the maximum dilution that prevented CPE formation. Additionally, neutralizing antibody titers against SARS-CoV-2 in the serum of monkeys that received three doses of BK2102 were quantified at day 42 with luciferase-expressing pseudovirus carrying the SARS-CoV-2 Wuhan strain spike (Abnova Corporation). Serum samples were 4-fold serially diluted and incubated at room temperature for 30 minutes with Spike-pseudovirus. Subsequently, the mixtures were added into wells containing 293-ACE2 cells. The luciferase expression of the infected cells was detected with Luciferase Assay System (Promega) after an incubation of 46 hours at 37 °C. NT50 neutralization titers were determined by a calibration curve with GraphPad prism.

### Enzyme-linked immune-sorbent assay (ELISA)

Half-well protein high-binding 96-well plates (Greiner) were coated with recombinant SARS-CoV-2 (D614G) spike or nucleocapsid proteins (SinoBiologicals) dissolved in PBS (50 ng/100 μL/well) and incubated at 4 °C overnight. 1% BSA PBS was used as blocking solution, and 1% BSA PBS-T was used to dilute the sera, antibodies, and streptavidin. Antigen-specific IgG antibodies in hamster sera were detected using horseradish peroxidase (HRP)-labeled goat anti-Syrian hamster IgG H+L Ab (Abcam, 1/30000 dilution). Biotinylated anti-Syrian hamster IgG1, IgG2/3 subclass Abs (Southern Biotech, 1/100 and 1/200000 dilutions, respectively) and IgA Ab (Brookwood Biomedical, 1/100 dilution) were used and detected with HRP-labeled streptavidin (Abcam, 1/1000 dilution). Mouse IgG was detected using HRP-labeled goat anti-mouse IgG H&L (Abcam, 1/100000 dilution). TMB one-component substrate and stopping solutions (Thermodics) were used for the chromogenic reaction, and the optical density (OD) was measured at 450-650 nm with a microplate reader. Endpoint titers of IgG were calculated based on a calibration curve using a five-parameter, non-linear curve fitting. Samples for which the endpoint titers could be calculated were defined as ‘Positive’, and the mean was calculated using only the titer of positive samples.

### Enzyme-linked immunosorbent spot (ELISPOT) assay

Hamster IFN-γ ELISPOT was performed using the ELISpot Flex: Hamster IFNLγ (ALP) (MABTECH) kit according to the manufacturer’s instructions. MSIP plates (Millipore) were washed 5 times with sterile water, coated with Capture mAb (MTH21) and incubated overnight at 4 °C. Coated plates were washed 5 times with PBS, blocked for 30Lminutes (at room temperature) with RPMI medium supplemented with 10% FBS and antibiotics. Two-hundred and fifty thousand splenocytes were seeded in each well and stimulated with SARS-CoV-2 spike or nucleocapsid peptide pools, consisting mainly of 15-mer sequences with 11-amino-acid (aa) overlaps (Miltenyi Biotec). Negative controls for non-stimulated cells were included in the test. After stimulation for a day at 37 °C, the spots were detected with the detection mAb (MTH29-biotin) and Streptavidin-ALP. After drying the plate, spots were counted using the ImmunoSpot^®^ S5 (Cellular Technology Limited). The average number of spots in the two negative control wells was subtracted from each well stimulated with peptide pools. The result was shown as the difference in spot-forming cells (SFC)/10^6^ splenocytes between the negative control and the peptide pool stimulated wells.

### mRNA and LNP production process

A codon-optimized mRNA encoding the SARS-CoV-2 S protein was *in vitro* synthesized and purified following the procedure of Moderna Therapeutics’ mRNA-1273 as reference (Hassett et al., 2019). The mRNA was encapsulated using a NanoAssemblr^TM^ Ignite^TM^ nanoparticle formulation system. The sample was subsequently concentrated to 1µg/100µL/dose using Amicon Ultra^®^ 4-100K, and filtered through a 0.45µm membrane for *in vivo* use.

### Evaluation of immunogenicity in hamsters

Four-week-old male Syrian hamsters were purchased from Japan SLC Inc. After a one-week housing period, they were first anesthetized via inhalation with 3% isoflurane, followed by intraperitoneal anesthesia with a combination of medetomidine, midazolam, and butorphanol (0.3, 4, and 5 mg/kg, respectively). One thousand or 1×10^4^ PFU of BK2102 were administered intra-nasally in a volume of 20 μL to confine administration to the upper respiratory tract. To evaluate the humoral immune response, blood samples were collected from the facial vein using a lancet (MEDIpoint), and neutralizing antibody titers in the sera were measured as described above. In addition, to evaluate mucosal immunity, nasal wash samples were collected through a plastic catheter using two separate washes of 1 mL of PBS. Only one animal inoculated BK2102 once was not used for IgA measurement because the nasal wash was contaminated with blood. To evaluate the cellular immune response, 1×10^4^ PFU of BK2102-inoculated hamsters were euthanized one week post-inoculation, and spleens were collected. The spleens were mechanically crushed with the piston of a syringe and passed through a cell strainer (100 μm). Cell suspensions were treated with RBC lysis buffer (Biolegend) to remove red blood cells, as per manufacturer instructions. The spleen cells were then suspended in RPMI medium supplemented with 10% FBS and antibiotics at a concentration of 1×10^6^ cells/mL and stimulated by SARS-CoV-2 spike or nucleocapsid peptide pools, consisting mainly of 15-mer sequences with 11-aa overlaps (Miltenyi Biotec). Negative controls for non-stimulated cells were included in the test. Stimulated cells were incubated at 37 °C for one day, and supernatants were collected and stored at -80 °C until use. Interferon-gamma (IFN-γ) and interleukin-4 (IL-4) were quantified with commercially-available ELISA kits (MABTECH AB and FineTest, respectively) following the manufacturer’s protocol. Furthermore, IFN-γ-producing cells were identified after a 22-hour stimulation culture with spike or nucleocapsid peptide pools, and analyzed by ELISpot Flex (MABTECH AB). We also performed a challenge assay to investigate whether these immune responses were effective in protecting against other variants. Four weeks post-inoculation with 1×10^3^ PFU or 1×10^4^ PFU of BK2102, 3×10^5^ PFU of SARS-CoV-2 D614G strain, TY41-702 strain (Omicron variant BA.5) or gamma strain were used to challenge the hamsters through the intra-nasal route to target the lower respiratory tract. The weights of the hamsters were monitored daily, and they were euthanized four days after the challenge. Lungs were divided, cut into small pieces, and homogenized with a biomasher II device in 500 μL of D-MEM. Supernatants were collected as lung homogenates after centrifugation at 300 ×g for 5 min at 4 °C. Nasal wash specimens were obtained by flushing one mL of D-PBS into the nasal cavity from the trachea in the direction of the nose where it was collected. Infectious virus titers in these samples were evaluated using a plaque formation assay, as described above. To evaluate immune response persistence, hamsters inoculated with 1×10^3^ PFU or 1×10^4^ PFU of BK2102 were maintained for 364 days post-infection. Blood samples were collected from the facial vein using a lancet (MEDIpoint), and neutralizing antibody titers were measured. At 420 days after inoculation, a viral challenge assay was performed to evaluate whether the immunity contributing to infection protection was maintained. Hamsters were infected with 3×10^5^ PFU of the SARS-CoV-2 D614G strain, and weight changes were monitored daily for 10 days post-infection.

### Evaluation of immunogenicity in monkeys

Sixteen male and female cynomolgus monkeys, two-to three years old, were purchased from Hamri Inc. After a one-week acclimatization housing period, they were sedated with a mixture of ketamine (5 mg/kg) and xylazine (2 mg/kg) administered intramuscularly. Ten million PFU of BK2102 were inoculated intranasally in a volume of 1.6 mL. Four monkeys were inoculated with a single-dose of BK2102, and blood samples were collected at 0, 28, 35 and 42 days after inoculation from the radial vein, femoral vein, or abdominal vena cava using a syringe. Neutralizing antibody titers in the sera were measured as described above to evaluate the humoral immune response. Another twelve monkeys were inoculated with 1×10^7^ PFU of BK2102 or the solvent, receiving three doses given at two-week intervals. Blood samples were collected 42 days after the last inoculation, and neutralizing antibody titers in the sera were measured.

### Evaluation of BK2102 pathogenicity in hamsters

Four-week-old male Syrian hamsters were obtained from Japan SLC Co. Ltd. After acclimatization, 1×10^4^ PFU of BK2102 or 1×10^4^ PFU of the SARS-CoV-2 D614G strain were used for infection, as described above. These and naïve hamsters were euthanized three days post-infection. The heads and lungs were collected and fixed in 10% formalin. Sections of the head were prepared to expose different regions of the nasal cavity and were stained with hematoxylin-eosin or an immunohistochemistry (IHC) staining kit using a SARS-CoV-2 spike RBD-specific antibody (Sino Biological). The damage score of each section was defined as 0: not remarkable (< 10%); 1: minimal (10-50%); and 2: mild (50-70%).

### *In vivo* passage of BK2102 in hamsters

Four-week-old male Syrian hamsters were obtained from Japan SLC Co. Ltd. After a one-week acclimatization period, 1×10^5^ PFU of BK2102 or A50-18 strain (a TS mutant isolated in a previous report) in a volume of 20 μL were used to infect the hamsters intranasally. The hamsters were observed daily, and their body weights were measured at 0 and 3 days post-infection. At this point, hamsters were euthanized, and nasal wash specimens were collected with 500 μL D-PBS. After filtration through 0.45-μm and 0.22-μm filters, 100 μL were used to infect a new group of naive hamsters. The passage of inoculum was repeated three times, and nasal wash samples were collected at every passage. The viral genome in these nasal wash samples was quantified via qPCR, as described above. Sanger sequencing was performed to analyze the mutations introduced to generate an attenuated phenotype.

### Generation of human ACE2-transgenic mice

The transgene was prepared as described previously (Ikawa et al., 1995; Okabe et al., 1997). Briefly, the hACE2-coding sequence was amplified via PCR with the following primers: 5′-aatctagagccgccgccgccatgtcaagctcttcctggctccttc-3′ and 5′-aaactcgagctaaaaggaggtctgaacatcatca-3′, using human testis cDNA as the template. The *Xba*I and *Xho*I sites included in the PCR primers were used to introduce the amplified hACE2 cDNA into a pCAG1.1 expression vector (https://www.addgene.org/173685/) containing the chicken *beta-actin* promoter and cytomegalovirus enhancer, the *beta-actin* intron, and the rabbit *globin* poly-adenylation signal. The transgene fragment was excised using *Sac*I and *Pac*I and gel-purified. Transgenic mouse lines were generated by injecting purified transgene fragments into C57BL/6N×C57BL/6N fertilized eggs. A total of 350 DNA-injected eggs were transplanted into pseudopregnant mice, resulting in 32 newborn pups. Three of these were transgenic, and the first line was established as Tg (CAG-hACE2)01Osb. The Tg mice were kept of a B6D2F1 background. Expression of hACE2 in each organ was confirmed via western blotting. Briefly, organ homogenates were prepared in radioimmunoprecipitation assay buffer containing a proteinase inhibitor cocktail (Thermo Scientific) using a bead mill homogenizer (Fischer Scientific). Protein concentrations were measured via BCA assay (Pierce), and 10 μg of protein were subjected to SDS-PAGE and subsequent western blotting. hACE2 and beta-actin were detected using a rabbit polyclonal antibody against hACE2 (Abcam: ab15348) and a mouse monoclonal antibody to beta-actin (Abcam: ab8226) with Can Get Signal solutions (TOYOBO).

### Evaluation of safety in transgenic mice

Seven- to nine-week-old Tg mice were anesthetized via inhalation of 3% isoflurane, then receiving a combination of medetomidine, midazolam, and butorphanol (0.3, 4, and 5 mg/kg, respectively) intraperitoneally. Mice were infected with SARS-CoV-2 in a volume of 20 μL, and their weights were measured daily. Mice that reached humane endpoints, such as difficulty walking or rapid weight loss, were euthanized by bleeding under isoflurane anesthesia. The brain, olfactory bulbs, nasal turbines, and lungs were collected. Mice infected with live-attenuated strains that did not reach the humane endpoint were also euthanized 14 days post-infection.

### Statistical analysis

Two-way analysis of variance (ANOVA), one-way ANOVA or Mann–Whitney *U* test was used to calculate statistical significance. These data were analyzed using GraphPad Prism 9.4.1 software. Statistical significance was set at a *p*-value < 0.05.

## ACKNOWLEDGMENTS

We appreciate the assistance of Paola Miyazato, Manae Morishima, and Kaori Yamamoto from The Research Foundation for Microbial Diseases of Osaka University (BIKEN). The authors thank Shiho Torii, Chikako Ono, and Yoshiharu Matsuura for their CPER technical support. We would like to thank Mitsuko Mori for generation of human ACE2-transgenic mice. The authors also acknowledge the NGS Core facility of the Genome Information Research Center at the Research Institute for Microbial Diseases of Osaka University for their support with next-generation sequencing analyses. This work was conducted as part of “The Research Foundation for Microbial Diseases of Osaka University Project for Infectious Disease Prevention”.

## AUTHOR CONTRIBUTIONS

M.S. and S.O. conducted and controlled most of the experiments and prepared the manuscript. W.K. performed the *in vivo* passage assays. T.N. and M.I. established Tg-mice. A.Y., and S.G. constructed various recombinant viruses. T.G., and H.S. supported the experiments related to prolonged protection and *in vivo* passage assays. L.S. conducted the cellular immunity assays. M.M. and T.M. measured spike-specific IgA titer by ELISA. M.Y. performed neutralizing antibody assay of monkeys using luciferase-expressing pseudovirus carrying the SARS-CoV-2 Wuhan strain’s spike protein. S.T., K.Y., and H.E. designed and managed the study.

## Supplementary figure legends

**Supplementary fig. 1 Characteristics of the vaccine candidates.**

(A) Candidate vaccine constructs. The abbreviations shown in the figure are respectively, ORF: Open Reading Frame, NSP: Non-Structural Protein, FCS: Furin Cleavage Site.

(B) Immunogenicity of vaccine candidates in hamsters. Neutralizing antibody titers in the sera were measured 21 days post-inoculation.

**Supplementary fig. 2 Comparison of immunogenicity of BK2102 with other vaccine modalities**

(A) An mRNA vaccine was administered to hamsters intramuscularly twice at 2-week intervals, and BK2102 was inoculated to another group of hamsters intranasally once or twice at 2-week intervals. Neutralizing antibodies in the sera were measured at day 28 post-first inoculation using authentic SARS-CoV-2 wild-type D614G strain. The LOD was 2^3^, and for samples below the LOD, the mean value was set to 2^2^. The dotted line represents the assay’s LOD. For statistical analysis, one-way ANOVA with Tukey’s multiple comparison test was performed (ns, not significant; *, *p* < 0.05; **, *p* < 0.01; ***, *p* < 0.001).

(B) SARS-CoV-2 spike-specific IgA measured via ELISA of nasal wash from inoculated hamsters, 28 days post-first inoculation. To calculate the IgA titer, the dilution factor at an OD value of 0.1 was considered and corrected with the value of the standard serum included in each plate. For statistical analysis, one-way ANOVA with Tukey’s multiple comparison test was performed (ns, not significant; *, *p* < 0.05; **, *p* < 0.01; ***, *p* < 0.001).

(C) SARS-CoV-2 spike-specific total IgG and IgG subclasses (IgG1 and IgG2/3) were measured in the serum of inoculated hamsters, 28 days post-first inoculation via ELISA.

(D) Recombinant SARS-CoV-2 Spike-protein mixed with an alum adjuvant was administered to hamsters intramuscularly twice at 2-week intervals, and BK2102 inoculated once to another group of hamsters intranasally. SARS-CoV-2 spike-specific total IgG and IgG subclasses (IgG1 and IgG2/3) were measured via ELISA of serum from inoculated hamsters, 28 days post-inoculation.

**Supplementary fig. 3 Immunogenicity of BK2102 in monkeys**

(A) Immune response in monkeys. Neutralizing antibodies in the sera of monkeys inoculated with 10^7^ PFU of BK2102 was measured at the indicated time points post-inoculation. The data for individual monkeys are shown. The LOD was 2^3^, and for samples below the LOD, the mean value was set to 2^2^. The dotted line represents the assay’s LOD.

(B) Neutralizing antibodies in the sera were induced in BK2102-inoculated monkeys. Monkeys were inoculated with 10^7^ PFU of BK2102 or the solvent, receiving three doses given at two-week intervals. Neutralizing antibodies in the sera were measured at day 42 post-inoculation using luciferase-expressing pseudovirus carrying the SARS-CoV-2 Wuhan strain’s spike protein. Symbols represent titers of individual animals, and the bars indicate the median. The LOD was 2^3.8^, and for samples below the LOD, the mean value was c set to 2^2^. The dotted line represents the assay’s LOD. For statistical analysis, Mann–Whitney *U* test was performed (**, *p* < 0.01).

**Supplementary fig. 4 Persistence of the neutralizing antibodies induced by BK2102 in each group.**

Results for individual mice in the group used in the experiment of Figure 2D are shown. Light-blue circles correspond to mice inoculated with 10^3^ PFU, while dark-blue circles correspond to mice inoculated with 10^4^ PFU. Dotted lines depict one dose, and solid lines to two doses. ¶ Hamsters died due to aging, fighting, or mishandling.

**Supplementary fig. 5 BK2102 induces protective immunity against SARS-CoV-2 gamma strain**

(A) Neutralizing antibodies in the sera were induced by BK2102-inoculated hamsters. Neutralizing antibodies in the sera were measured at day 28 post-inoculation using the following authentic SARS-CoV-2 strains: wild-type D614G (left) and gamma (right). Symbols represent titers of individual animals, and the bars indicate the median. The dotted line represents the assay’s LOD. For statistical analysis, one-way ANOVA with Tukey’s multiple comparison test was performed (ns, not significant; ****, *p* < 0.0001).

(B) Weight changes in vaccinated mice after the challenge with the gamma strain. Hamsters inoculated with BK2102 were challenged with 3×10^5^ PFU of the gamma strain on day 28 post-inoculation. The symbols represent the average weight of the hamsters, and error bars indicate the mean SD. Two-way ANOVA with Tukey’s multiple comparison test was performed for statistical analysis (****, *p* < 0.0001).

**Supplementary fig. 6 Evaluation of BK2102 onward transmission in hamsters**

(A) Scheme of the onward transmission experiment. Hamsters that received the full vaccination protocol with the indicated doses of BK2102 and recombinant SARS-CoV-2 spike protein mixed with an alum adjuvant were challenged with the wild-type D614G strain. Vaccinated and infected animals were co-housed with naïve hamsters at one day post-infection.

(B) Body weight was monitored for six days.

**Supplementary fig. 7 Evaluation of the tissue damage induced by BK2102**

H&E staining of nasal cavity sections and lungs is shown for hamsters infected with the wild-type D614G strain or the BK2102 vaccine candidate. IHC results using a SARS-CoV-2 spike RBD-specific antibody are shown in for the same sections. Scale bar: 200 μm.

**Supplementary fig. 8 BK2102 showed a low risk of transmission**

The body weights of hamsters inoculated with the A50-18 TS and BK2102 strains obtained in the *in vivo* passage experiment detailed in Figure 4D are shown. For statistical analysis, two-way ANOVA with Holm-Sidak’s multiple comparison test was performed (ns, not significant; **, p < 0.01; *** p < 0.001; **** p < 0.0001).

**Supplementary fig. 9 Expression of hACE2 and body weight of Tg mice after infection**

(A) The expression of hACE2 in various tissues of Tg mice was detected via western blotting using an anti-human ACE2 antibody (ab15348).

(B) The body weight of Tg mice infected with the wild-type D614G, B-1 ΔFCS, L50-33, and A50-18 TS strains. The symbols represent the average weight of each group, and error bars indicate the mean SD. Two-way ANOVA with Tukey’s multiple comparison test was performed for statistical analysis (ns, not significant; ****, *p* < 0.0001).

(C) The body weight of Tg mice inoculated with the BK2102. The symbols represent the average weight of each group, and error bars indicate the mean SD. Two-way ANOVA with Tukey’s multiple comparison test was performed for statistical analysis (ns, not significant).

## References

Asaka, M.N., Utsumi, D., Kamada, H., Nagata, S., Nakachi, Y., Yamaguchi, T., Kawaoka, Y., Kuba, K., and Yasutomi, Y. (2021). Highly susceptible SARS-CoV-2 model in CAG promoter-driven hACE2-transgenic mice. JCI Insight 6. 10.1172/jci.insight.152529.

Baden, L.R., El Sahly, H.M., Essink, B., Kotloff, K., Frey, S., Novak, R., Diemert, D., Spector, S.A., Rouphael, N., Creech, C.B., et al. (2021). Efficacy and Safety of the mRNA-1273 SARS-CoV-2 Vaccine. N Engl J Med 384, 403–416. 10.1056/NEJMoa2035389.

Bao, L., Deng, W., Huang, B., Gao, H., Liu, J., Ren, L., Wei, Q., Yu, P., Xu, Y., Qi, F., et al. (2020). The pathogenicity of SARS-CoV-2 in hACE2 transgenic mice. Nature 583, 830–833. 10.1038/s41586-020-2312-y.

Bull, J.J. (2015). Evolutionary reversion of live viral vaccines: Can genetic engineering subdue it? Virus Evol 1. 10.1093/ve/vev005.

Chiuppesi, F., Nguyen, V.H., Park, Y., Contreras, H., Karpinski, V., Faircloth, K., Nguyen, J., Kha, M., Johnson, D., Martinez, J., et al. (2022). Synthetic multiantigen MVA vaccine COH04S1 protects against SARS-CoV-2 in Syrian hamsters and non-human primates. NPJ Vaccines 7, 7. 10.1038/s41541-022-00436-6.

Cong, Y., Ulasli, M., Schepers, H., Mauthe, M., V’Kovski, P., Kriegenburg, F., Thiel, V., de Haan, C.A.M., and Reggiori, F. (2020). Nucleocapsid Protein Recruitment to Replication-Transcription Complexes Plays a Crucial Role in Coronaviral Life Cycle. J Virol 94. 10.1128/JVI.01925-19.

Davidson, A.D., Williamson, M.K., Lewis, S., Shoemark, D., Carroll, M.W., Heesom, K.J., Zambon, M., Ellis, J., Lewis, P.A., Hiscox, J.A., and Matthews, D.A. (2020). Characterisation of the transcriptome and proteome of SARS-CoV-2 reveals a cell passage induced in-frame deletion of the furin-like cleavage site from the spike glycoprotein. Genome Med 12, 68. 10.1186/s13073-020-00763-0.

Desai, A., Shankar, S.K., Ravi, V., Chandramuki, A., and Gourie-Devi, M. (1995). Japanese encephalitis virus antigen in the human brain and its topographic distribution. Acta Neuropathol 89, 368–373. 10.1007/BF00309631.

Dutta, N.K., Mazumdar, K., and Gordy, J.T. (2020). The Nucleocapsid Protein of SARS-CoV-2: a Target for Vaccine Development. J Virol 94. 10.1128/JVI.00647-20.

Hasanpourghadi, M., Novikov, M., Ambrose, R., Chekaoui, A., Newman, D., Ding, J., Giles-Davis, W., Xiang, Z., Zhou, X.Y., Liu, Q., et al. (2023). Heterologous chimpanzee adenovirus vector immunizations for SARS-CoV-2 spike and nucleocapsid protect hamsters against COVID-19. Microbes Infect 25, 105082. 10.1016/j.micinf.2022.105082.

Hassett, K.J., Benenato, K.E., Jacquinet, E., Lee, A., Woods, A., Yuzhakov, O., Himansu, S., Deterling, J., Geilich, B.M., Ketova, T., et al. (2019). Optimization of Lipid Nanoparticles for Intramuscular Administration of mRNA Vaccines. Mol Ther Nucleic Acids 15, 1–11. 10.1016/j.omtn.2019.01.013.

Hoffmann, M., Kleine-Weber, H., and Pohlmann, S. (2020a). A Multibasic Cleavage Site in the Spike Protein of SARS-CoV-2 Is Essential for Infection of Human Lung Cells. Mol Cell 78, 779–784 e775. 10.1016/j.molcel.2020.04.022.

Hoffmann, M., Kleine-Weber, H., Schroeder, S., Kruger, N., Herrler, T., Erichsen, S., Schiergens, T.S., Herrler, G., Wu, N.H., Nitsche, A., et al. (2020b). SARS-CoV-2 Cell Entry Depends on ACE2 and TMPRSS2 and Is Blocked by a Clinically Proven Protease Inhibitor. Cell 181, 271–280 e278. 10.1016/j.cell.2020.02.052.

Hoft, D.F., Lottenbach, K.R., Blazevic, A., Turan, A., Blevins, T.P., Pacatte, T.P., Yu, Y., Mitchell, M.C., Hoft, S.G., and Belshe, R.B. (2017). Comparisons of the Humoral and Cellular Immune Responses Induced by Live Attenuated Influenza Vaccine and Inactivated Influenza Vaccine in Adults. Clin Vaccine Immunol 24. 10.1128/CVI.00414-16.

Ikawa, M., Kominami, K., Yoshimura, Y., Tanaka, K., Nishimune, Y., and Okabe, M. (1995). A rapid and non-invasive selection of transgenic embryos before implantation using green fluorescent protein (GFP). FEBS Lett 375, 125–128. 10.1016/0014-5793(95)01162-8.

Jha, N.K., Ojha, S., Jha, S.K., Dureja, H., Singh, S.K., Shukla, S.D., Chellappan, D.K., Gupta, G., Bhardwaj, S., Kumar, N., et al. (2021). Evidence of Coronavirus (CoV) Pathogenesis and Emerging Pathogen SARS-CoV-2 in the Nervous System: A Review on Neurological Impairments and Manifestations. J Mol Neurosci 71, 2192–2209. 10.1007/s12031-020-01767-6.

Johnson, B.A., Xie, X., Bailey, A.L., Kalveram, B., Lokugamage, K.G., Muruato, A., Zou, J., Zhang, X., Juelich, T., Smith, J.K., et al. (2021). Loss of furin cleavage site attenuates SARS-CoV-2 pathogenesis. Nature 591, 293–299. 10.1038/s41586-021-03237-4.

Johnston, S.C., Ricks, K.M., Jay, A., Raymond, J.L., Rossi, F., Zeng, X., Scruggs, J., Dyer, D., Frick, O., Koehler, J.W., et al. (2021). Development of a coronavirus disease 2019 nonhuman primate model using airborne exposure. PLoS One 16, e0246366. 10.1371/journal.pone.0246366.

Kumar, A., Pareek, V., Prasoon, P., Faiq, M.A., Kumar, P., Kumari, C., and Narayan, R.K. (2020). Possible routes of SARS-CoV-2 invasion in brain: In context of neurological symptoms in COVID-19 patients. J Neurosci Res 98, 2376–2383. 10.1002/jnr.24717.

Kushawaha, P.K., Gupta, R., Sundar, S., Sahasrabuddhe, A.A., and Dube, A. (2011). Elongation factor-2, a Th1 stimulatory protein of Leishmania donovani, generates strong IFN-gamma and IL-12 response in cured Leishmania-infected patients/hamsters and protects hamsters against Leishmania challenge. J Immunol 187, 6417–6427. 10.4049/jimmunol.1102081.

Lau, S.Y., Wang, P., Mok, B.W., Zhang, A.J., Chu, H., Lee, A.C., Deng, S., Chen, P., Chan, K.H., Song, W., et al. (2020). Attenuated SARS-CoV-2 variants with deletions at the S1/S2 junction. Emerg Microbes Infect 9, 837–842. 10.1080/22221751.2020.1756700.

Lin, J.W., Tang, C., Wei, H.C., Du, B., Chen, C., Wang, M., Zhou, Y., Yu, M.X., Cheng, L., Kuivanen, S., et al. (2021). Genomic monitoring of SARS-CoV-2 uncovers an Nsp1 deletion variant that modulates type I interferon response. Cell Host Microbe 29, 489–502 e488. 10.1016/j.chom.2021.01.015.

Lindeboom, R.G.H., Worlock, K.B., Dratva, L.M., Yoshida, M., Scobie, D., Wagstaffe, H.R., Richardson, L., Wilbrey-Clark, A., Barnes, J.L., Kretschmer, L., et al. (2024). Human SARS-CoV-2 challenge uncovers local and systemic response dynamics. Nature 631, 189–198. 10.1038/s41586-024-07575-x.

Machado, R.R.G., Walker, J.L., Scharton, D., Rafael, G.H., Mitchell, B.M., Reyna, R.A., de Souza, W.M., Liu, J., Walker, D.H., Plante, J.A., et al. (2023). Immunogenicity and efficacy of vaccine boosters against SARS-CoV-2 Omicron subvariant BA.5 in male Syrian hamsters. Nat Commun 14, 4260. 10.1038/s41467-023-40033-2.

Macklin, G.R., O’Reilly, K.M., Grassly, N.C., Edmunds, W.J., Mach, O., Santhana Gopala Krishnan, R., Voorman, A., Vertefeuille, J.F., Abdelwahab, J., Gumede, N., et al. (2020). Evolving epidemiology of poliovirus serotype 2 following withdrawal of the serotype 2 oral poliovirus vaccine. Science 368, 401–405. 10.1126/science.aba1238.

Marrack, P., McKee, A.S., and Munks, M.W. (2009). Towards an understanding of the adjuvant action of aluminium. Nat Rev Immunol 9, 287–293. 10.1038/nri2510.

Natekar, J.P., Pathak, H., Stone, S., Kumari, P., Sharma, S., Auroni, T.T., Arora, K., Rothan, H.A., and Kumar, M. (2022). Differential Pathogenesis of SARS-CoV-2 Variants of Concern in Human ACE2-Expressing Mice. Viruses 14. 10.3390/v14061139.

Nouailles, G., Adler, J.M., Pennitz, P., Peidli, S., Teixeira Alves, L.G., Baumgardt, M., Bushe, J., Voss, A., Langenhagen, A., Langner, C., et al. (2023). Live-attenuated vaccine sCPD9 elicits superior mucosal and systemic immunity to SARS-CoV-2 variants in hamsters. Nat Microbiol 8, 860–874. 10.1038/s41564-023-01352-8.

Okabe, M., Ikawa, M., Kominami, K., Nakanishi, T., and Nishimune, Y. (1997). ’Green mice’ as a source of ubiquitous green cells. FEBS Lett 407, 313–319. 10.1016/s0014-5793(97)00313-x.

Okamura, S., Yoshida, A., Miyazato, P., Matsumoto, M., and Ebina, H. (2023). Protocol to isolate temperature-sensitive SARS-CoV-2 mutants and identify associated mutations. STAR Protoc 4, 102352. 10.1016/j.xpro.2023.102352.

Peacock, T.P., Goldhill, D.H., Zhou, J., Baillon, L., Frise, R., Swann, O.C., Kugathasan, R., Penn, R., Brown, J.C., Sanchez-David, R.Y., et al. (2021). The furin cleavage site in the SARS-CoV-2 spike protein is required for transmission in ferrets. Nat Microbiol 6, 899–909. 10.1038/s41564-021-00908-w.

Ploquin, A., Szecsi, J., Mathieu, C., Guillaume, V., Barateau, V., Ong, K.C., Wong, K.T., Cosset, F.L., Horvat, B., and Salvetti, A. (2013). Protection against henipavirus infection by use of recombinant adeno-associated virus-vector vaccines. J Infect Dis 207, 469–478. 10.1093/infdis/jis699.

Polack, F.P., Thomas, S.J., Kitchin, N., Absalon, J., Gurtman, A., Lockhart, S., Perez, J.L., Perez Marc, G., Moreira, E.D., Zerbini, C., et al. (2020). Safety and Efficacy of the BNT162b2 mRNA Covid-19 Vaccine. N Engl J Med 383, 2603–2615. 10.1056/NEJMoa2034577.

Primard, C., Monchatre-Leroy, E., Del Campo, J., Valsesia, S., Nikly, E., Chevandier, M., Boue, F., Servat, A., Wasniewski, M., Picard-Meyer, E., et al. (2023). OVX033, a nucleocapsid-based vaccine candidate, provides broad-spectrum protection against SARS-CoV-2 variants in a hamster challenge model. Front Immunol 14, 1188605. 10.3389/fimmu.2023.1188605.

Rosenke, K., Meade-White, K., Letko, M., Clancy, C., Hansen, F., Liu, Y., Okumura, A., Tang-Huau, T.L., Li, R., Saturday, G., et al. (2020). Defining the Syrian hamster as a highly susceptible preclinical model for SARS-CoV-2 infection. Emerg Microbes Infect 9, 2673–2684. 10.1080/22221751.2020.1858177.

Sabin, A.B. (1985). Oral poliovirus vaccine: history of its development and use and current challenge to eliminate poliomyelitis from the world. J Infect Dis 151, 420–436. 10.1093/infdis/151.3.420.

Sasaki, M., Toba, S., Itakura, Y., Chambaro, H.M., Kishimoto, M., Tabata, K., Intaruck, K., Uemura, K., Sanaki, T., Sato, A., et al. (2021). SARS-CoV-2 Bearing a Mutation at the S1/S2 Cleavage Site Exhibits Attenuated Virulence and Confers Protective Immunity. mBio 12, e0141521. 10.1128/mBio.01415-21.

Seo, S.H., and Jang, Y. (2020). Cold-Adapted Live Attenuated SARS-Cov-2 Vaccine Completely Protects Human ACE2 Transgenic Mice from SARS-Cov-2 Infection. Vaccines (Basel) 8. 10.3390/vaccines8040584.

Thwaites, R.S., Uruchurtu, A.S.S., Negri, V.A., Cole, M.E., Singh, N., Poshai, N., Jackson, D., Hoschler, K., Baker, T., Scott, I.C., et al. (2023). Early mucosal events promote distinct mucosal and systemic antibody responses to live attenuated influenza vaccine. Nat Commun 14, 8053. 10.1038/s41467-023-43842-7.

Torii, S., Ono, C., Suzuki, R., Morioka, Y., Anzai, I., Fauzyah, Y., Maeda, Y., Kamitani, W., Fukuhara, T., and Matsuura, Y. (2021). Establishment of a reverse genetics system for SARS-CoV-2 using circular polymerase extension reaction. Cell Rep 35, 109014. 10.1016/j.celrep.2021.109014.

Trimpert, J., Dietert, K., Firsching, T.C., Ebert, N., Thi Nhu Thao, T., Vladimirova, D., Kaufer, S., Labroussaa, F., Abdelgawad, A., Conradie, A., et al. (2021). Development of safe and highly protective live-attenuated SARS-CoV-2 vaccine candidates by genome recoding. Cell Rep 36, 109493. 10.1016/j.celrep.2021.109493.

Ueno, S., Amarbayasgalan, S., Sugiura, Y., Takahashi, T., Shimizu, K., Nakagawa, K., Kawabata-Iwakawa, R., and Kamitani, W. (2024). Eight-amino-acid sequence at the N-terminus of SARS-CoV-2 nsp1 is involved in stabilizing viral genome replication. Virology 595. ARTN 110068 10.1016/j.virol.2024.110068.

Wang, P., Lau, S.Y., Deng, S., Chen, P., Mok, B.W., Zhang, A.J., Lee, A.C., Chan, K.H., Tam, R.C., Xu, H., et al. (2021a). Characterization of an attenuated SARS-CoV-2 variant with a deletion at the S1/S2 junction of the spike protein. Nat Commun 12, 2790. 10.1038/s41467-021-23166-0.

Wang, Y., Yang, C., Song, Y., Coleman, J.R., Stawowczyk, M., Tafrova, J., Tasker, S., Boltz, D., Baker, R., Garcia, L., et al. (2021b). Scalable live-attenuated SARS-CoV-2 vaccine candidate demonstrates preclinical safety and efficacy. Proc Natl Acad Sci U S A 118. 10.1073/pnas.2102775118.

Weinheimer, V.K., Becher, A., Tonnies, M., Holland, G., Knepper, J., Bauer, T.T., Schneider, P., Neudecker, J., Ruckert, J.C., Szymanski, K., et al. (2012). Influenza A viruses target type II pneumocytes in the human lung. J Infect Dis 206, 1685–1694. 10.1093/infdis/jis455.

Yasmin, F., Najeeb, H., Naeem, U., Moeed, A., Atif, A.R., Asghar, M.S., Nimri, N., Saleem, M., Bandyopadhyay, D., Krittanawong, C., et al. (2023). Adverse events following COVID-19 mRNA vaccines: A systematic review of cardiovascular complication, thrombosis, and thrombocytopenia. Immun Inflamm Dis 11, e807. 10.1002/iid3.807.

Yeh, M.T., Bujaki, E., Dolan, P.T., Smith, M., Wahid, R., Konz, J., Weiner, A.J., Bandyopadhyay, A.S., Van Damme, P., De Coster, I., et al. (2020). Engineering the Live-Attenuated Polio Vaccine to Prevent Reversion to Virulence. Cell Host Microbe 27, 736–751 e738. 10.1016/j.chom.2020.04.003.

Yeh, M.T., Smith, M., Carlyle, S., Konopka-Anstadt, J.L., Burns, C.C., Konz, J., Andino, R., and Macadam, A. (2023). Genetic stabilization of attenuated oral vaccines against poliovirus types 1 and 3. Nature 619, 135–142. 10.1038/s41586-023-06212-3.

Yoshida, A., Okamura, S., Torii, S., Komatsu, S., Miyazato, P., Sasaki, H., Ueno, S., Suzuki, H., Kamitani, W., Ono, C., et al. (2022). Versatile live-attenuated SARS-CoV-2 vaccine platform applicable to variants induces protective immunity. iScience 25, 105412. 10.1016/j.isci.2022.105412.

Young, B.E., Fong, S.W., Chan, Y.H., Mak, T.M., Ang, L.W., Anderson, D.E., Lee, C.Y., Amrun, S.N., Lee, B., Goh, Y.S., et al. (2020). Effects of a major deletion in the SARS-CoV-2 genome on the severity of infection and the inflammatory response: an observational cohort study. Lancet 396, 603–611. 10.1016/S0140-6736(20)31757-8.

Zhang, G.F., Meng, W., Chen, L., Ding, L., Feng, J., Perez, J., Ali, A., Sun, S., Liu, Z., Huang, Y., et al. (2022). Neutralizing antibodies to SARS-CoV-2 variants of concern including Delta and Omicron in subjects receiving mRNA-1273, BNT162b2, and Ad26.COV2.S vaccines. J Med Virol 94, 5678-5690. 10.1002/jmv.28032.

Zinzula, L. (2021). Lost in deletion: The enigmatic ORF8 protein of SARS-CoV-2. Biochem Biophys Res Commun 538, 116–124. 10.1016/j.bbrc.2020.10.045.

