## Supplementary figures and images for "Immunogenicity and safety of a live-attenuated SARS-CoV-2 vaccine candidate based on multiple attenuation mechanisms"

### Supplemental fig 1

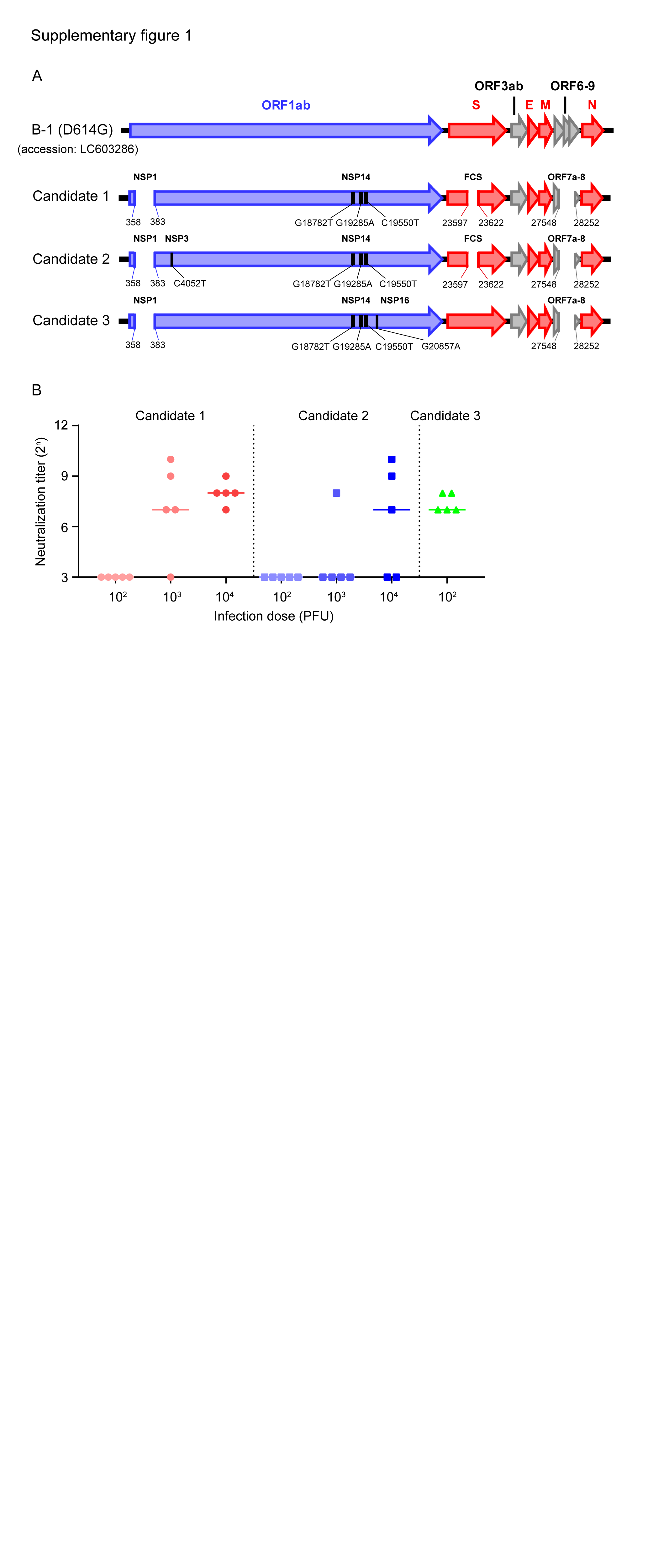

### Supplemental fig 2

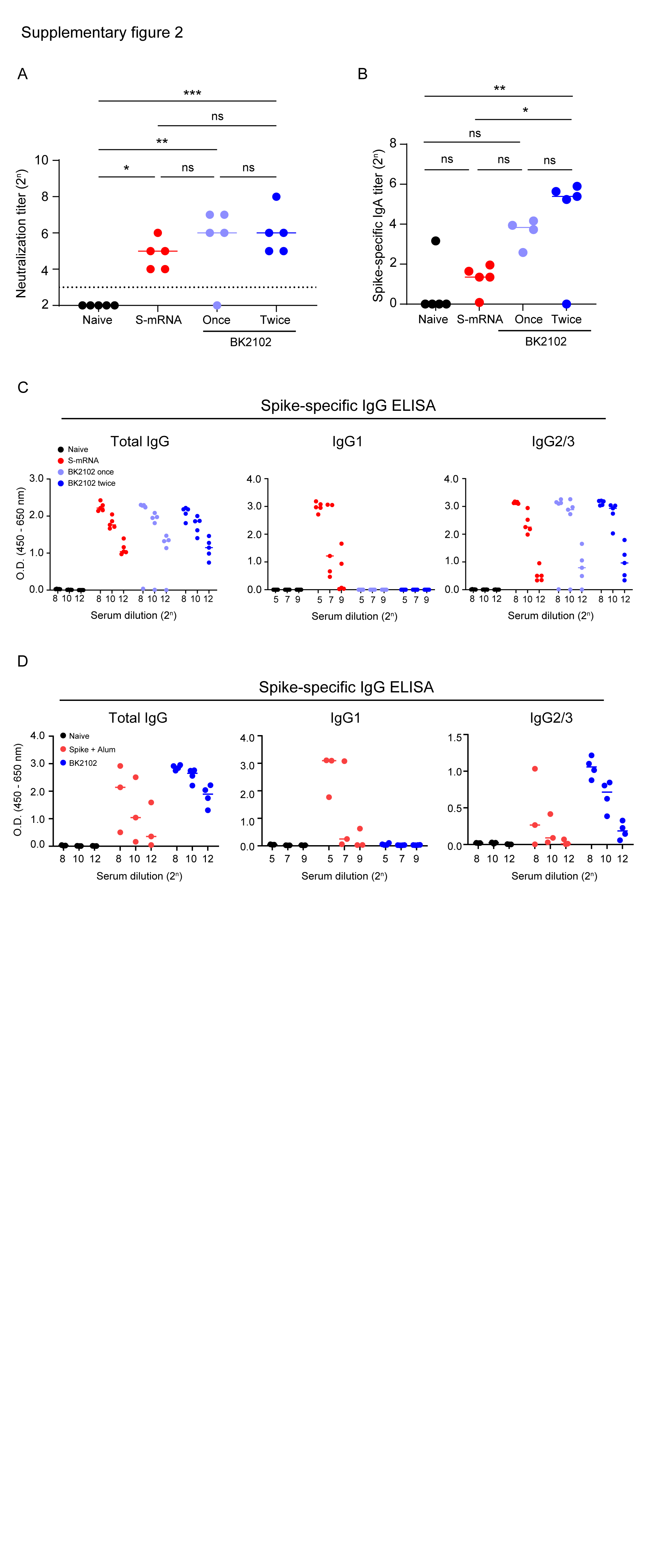

### Supplemental fig 3

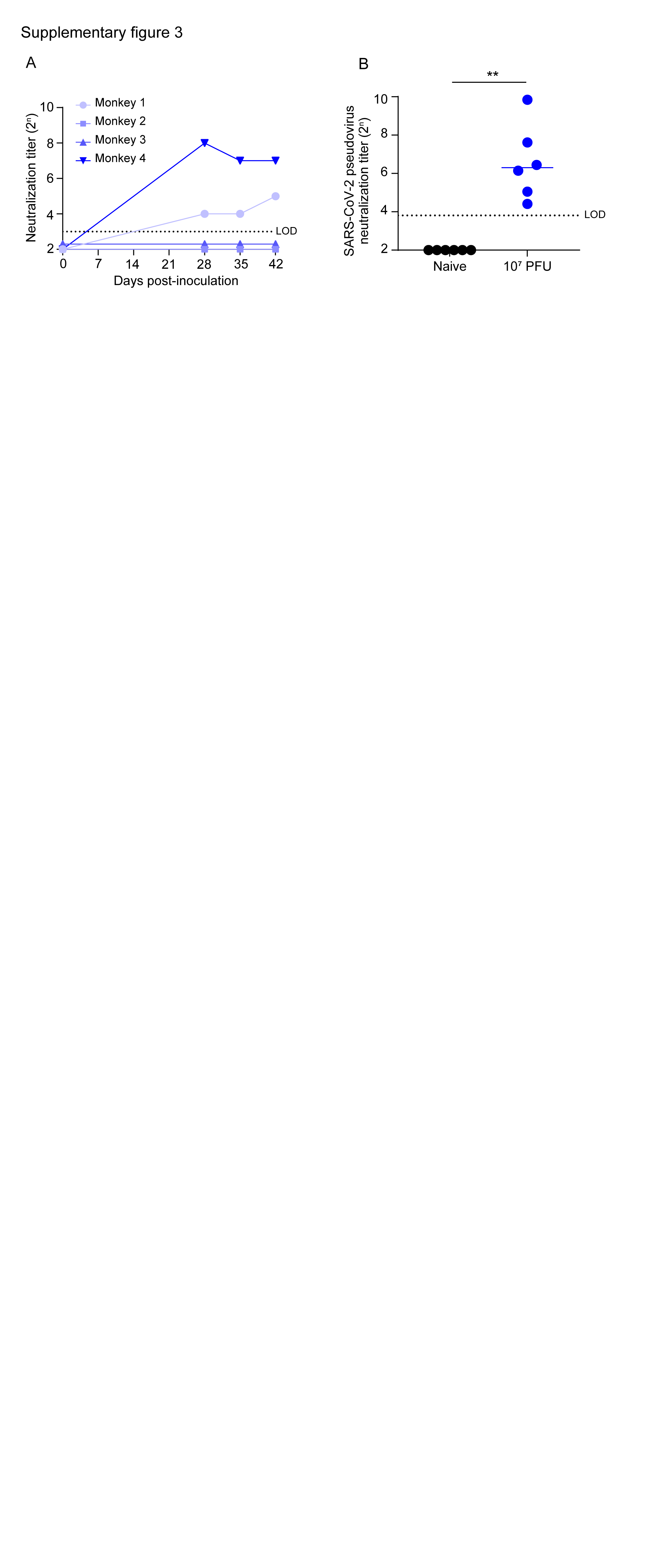

### Supplemental fig 4

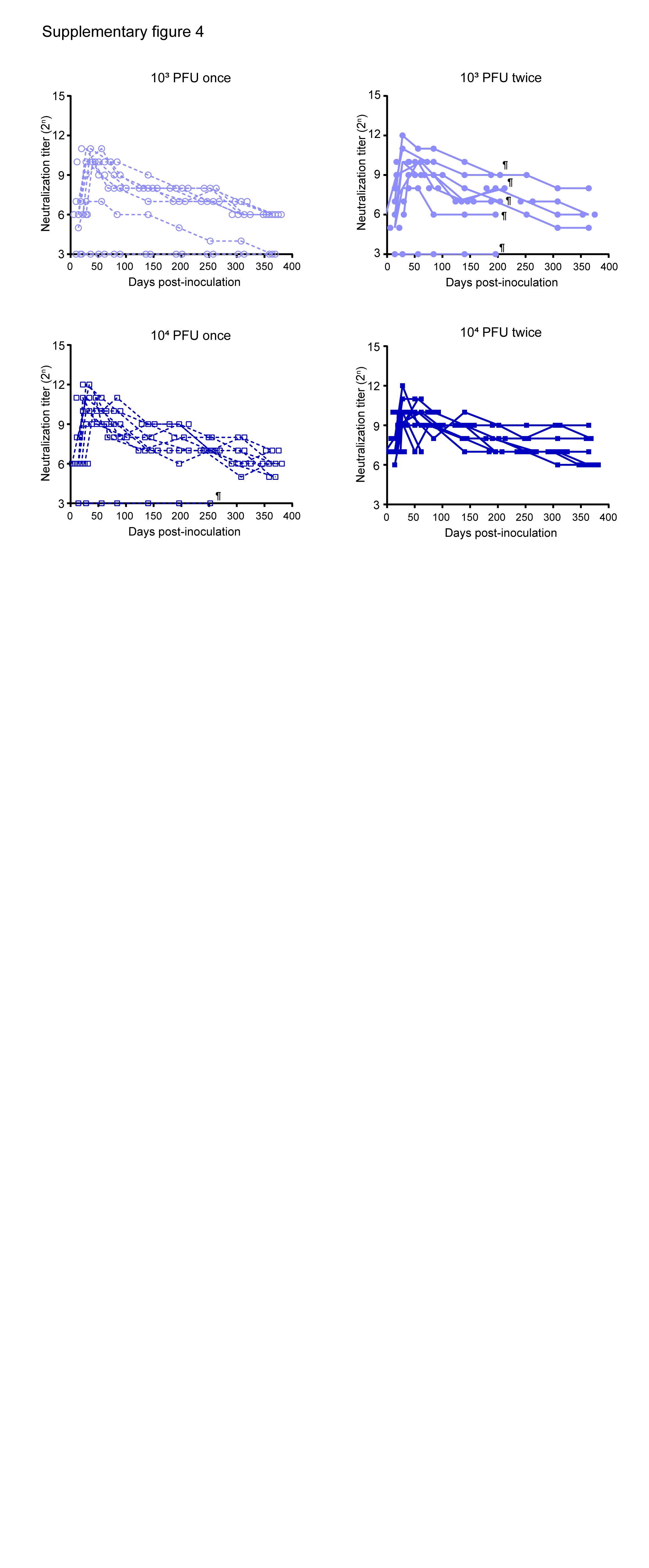

### Supplemental fig 5

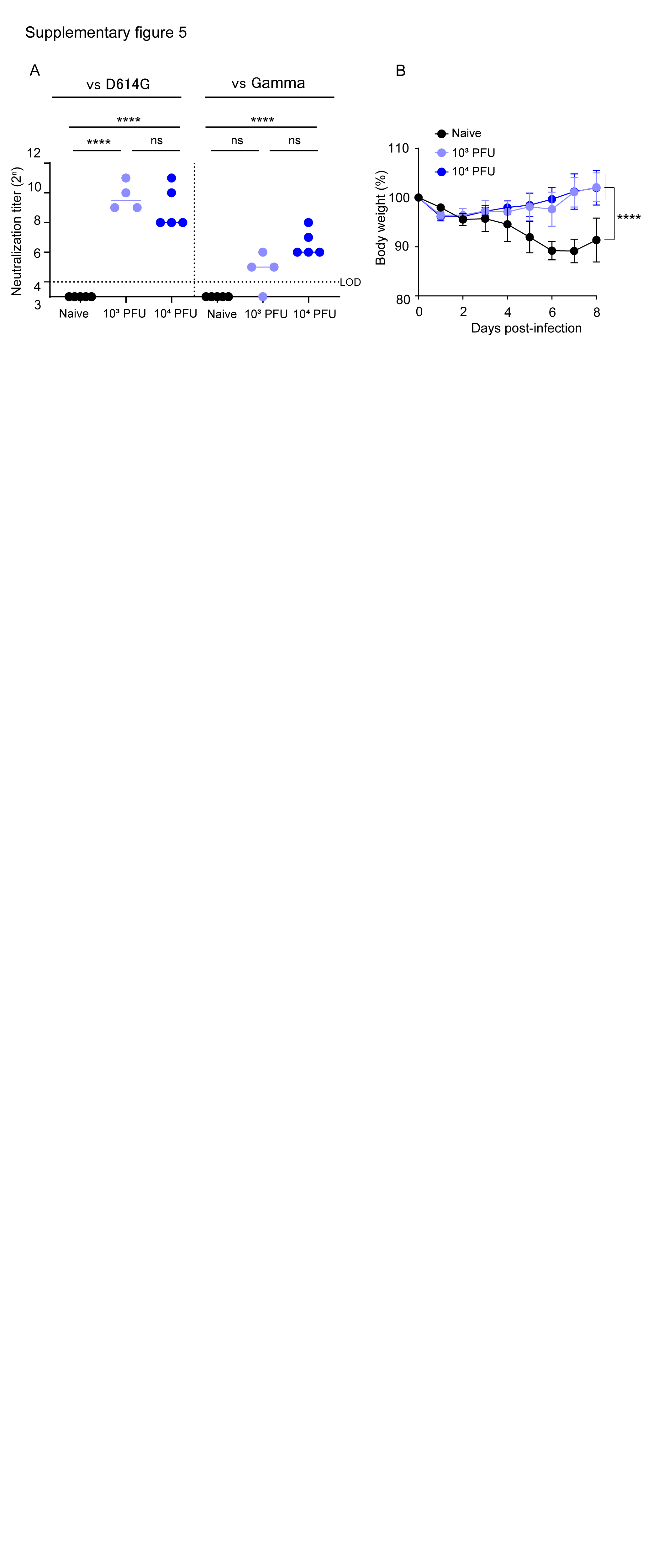

### Supplemental fig 6

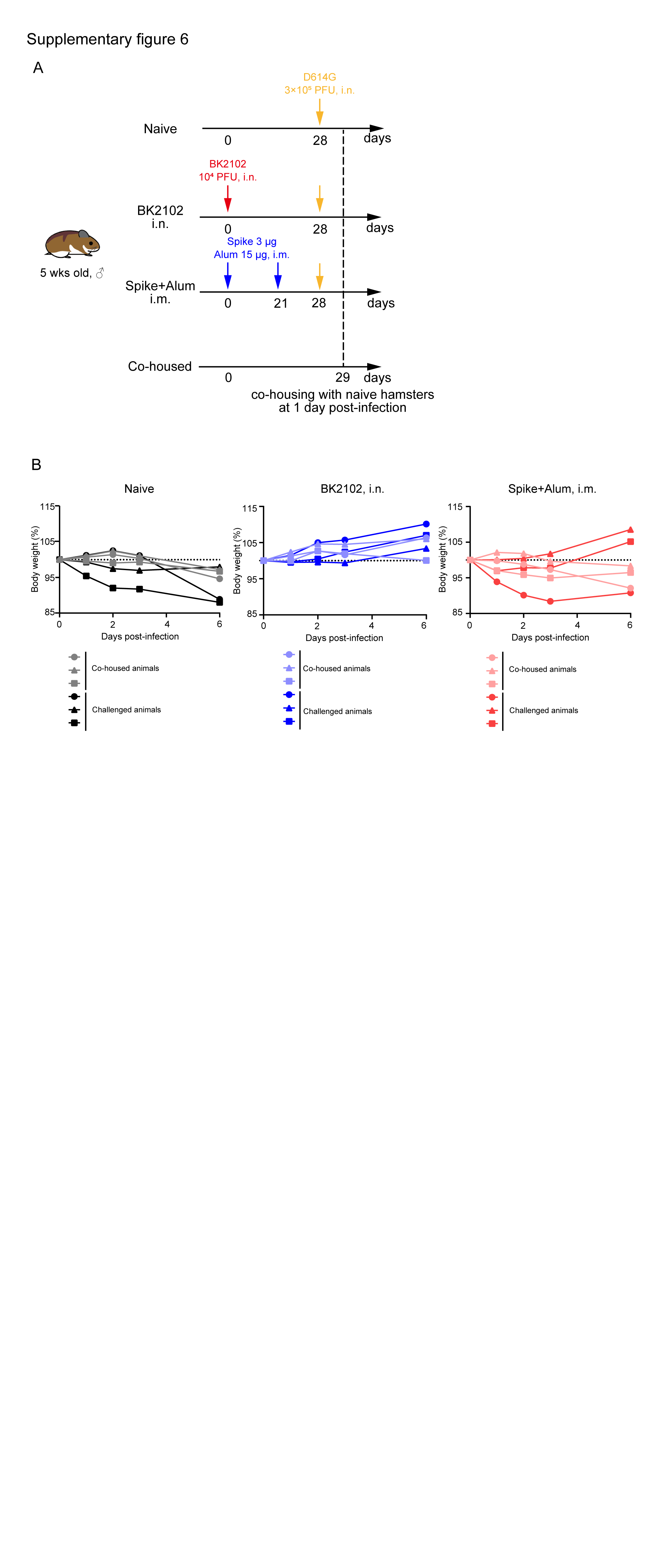

### Supplemental fig 7

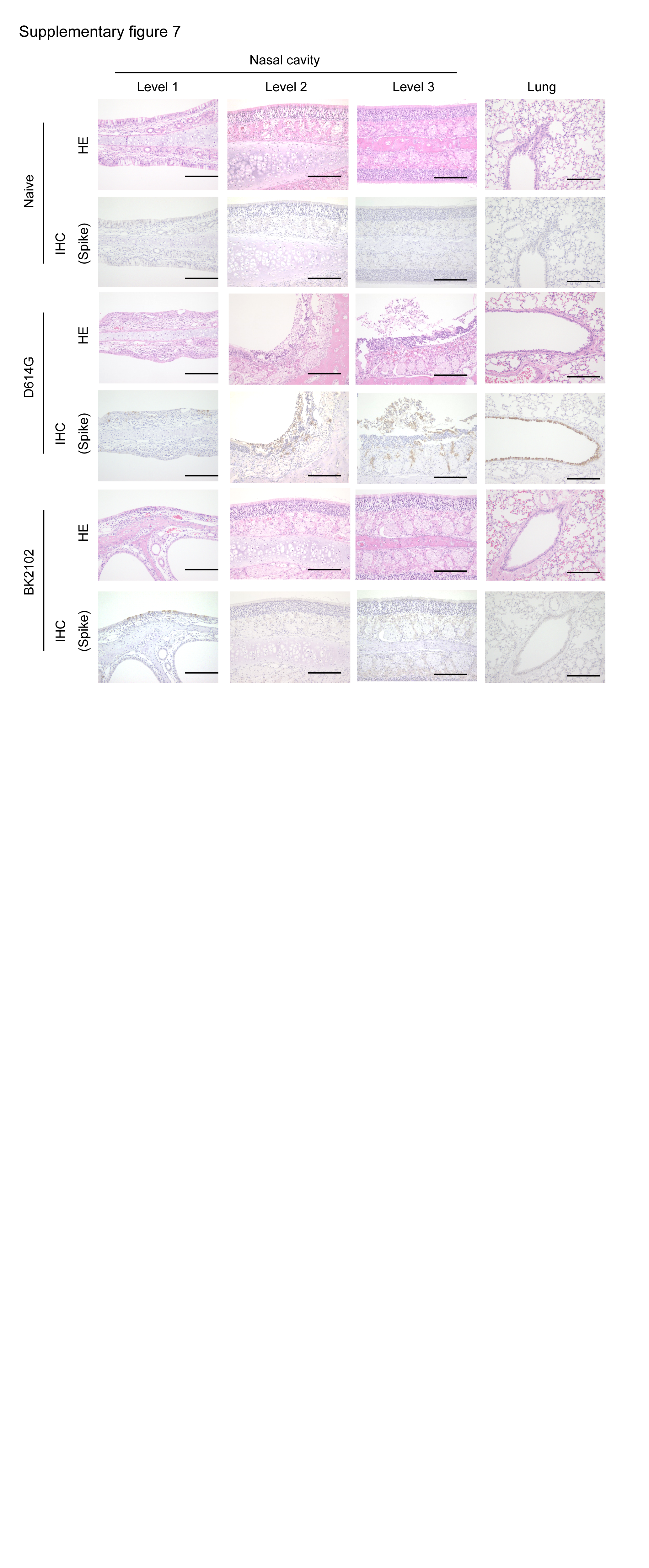

### Supplemental fig 8

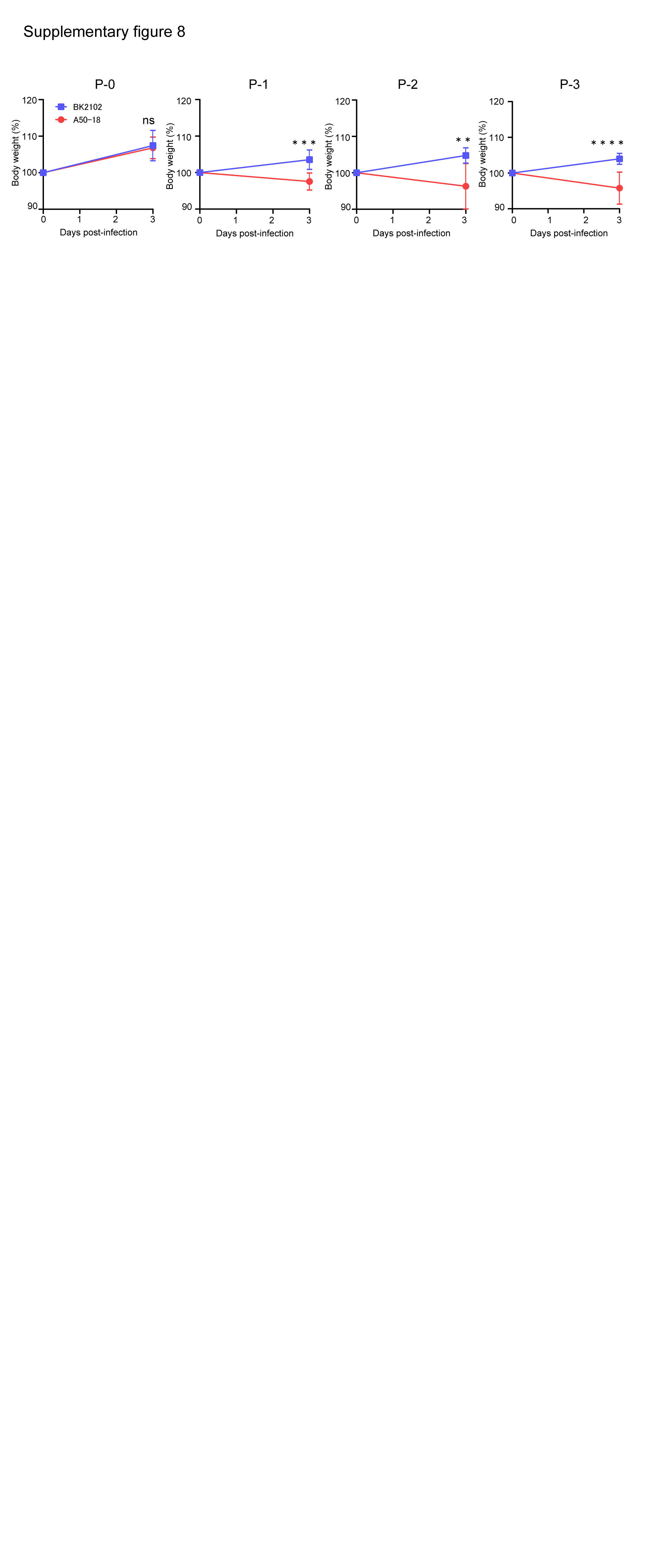

### Supplemental fig 9

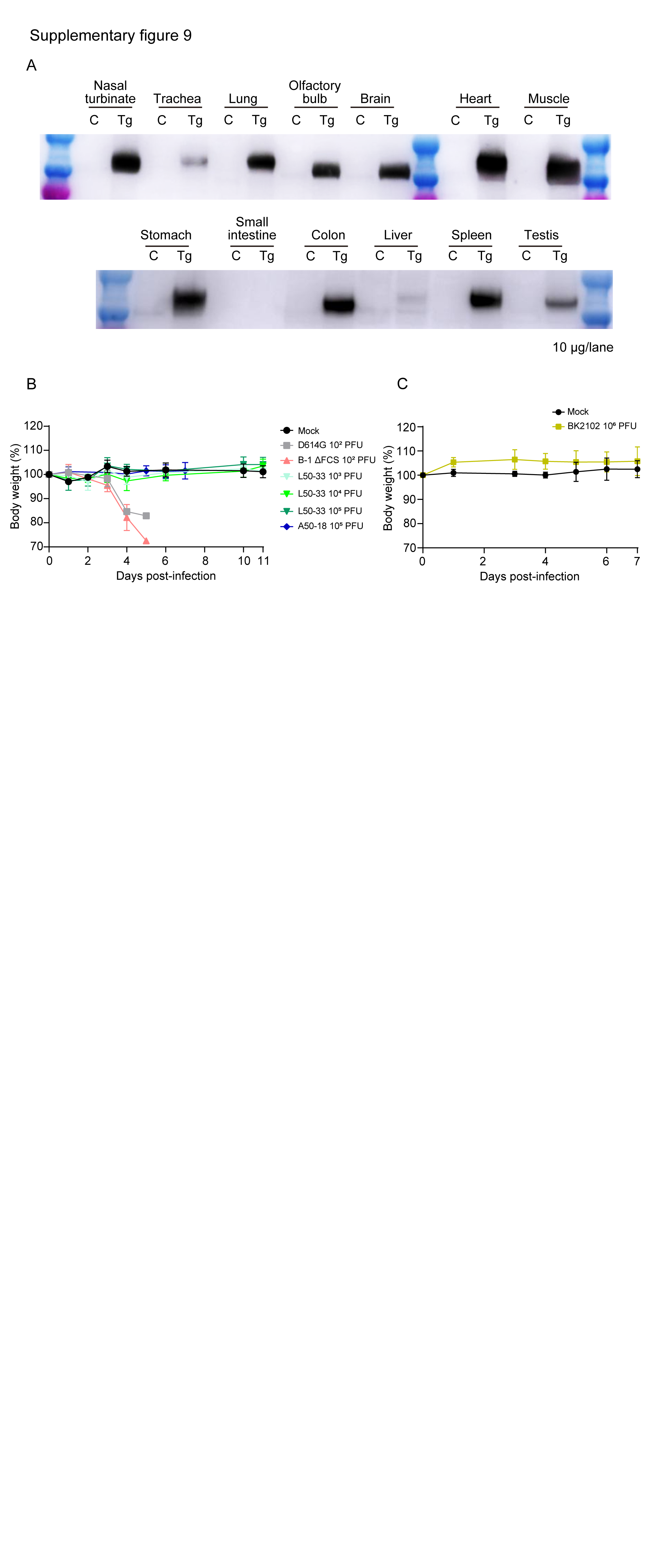
